# Tree Lab: Portable genomics for early detection of plant viruses and pests in Sub-Saharan Africa

**DOI:** 10.1101/702613

**Authors:** Laura M. Boykin, Peter Sseruwagi, Titus Alicai, Elijah Ateka, Ibrahim Umar Mohammed, Jo-Ann L. Stanton, Charles Kayuki, Deogratius Mark, Tarcisius Fute, Joel Erasto, Hilda Bachwenkizi, Brenda Muga, Naomi Mumo, Jenniffer Mwangi, Phillip Abidrabo, Geoffrey Okao-Okuja, Geresemu Omuut, Jacinta Akol, Hellen B. Apio, Francis Osingada, Monica A. Kehoe, David Eccles, Anders Savill, Stephen Lamb, Tonny Kinene, Christopher B. Rawle, Abishek Muralidhar, Kirsty Mayall, Fred Tairo, Joseph Ndunguru

## Abstract

In this case study we successfully teamed the PDQeX DNA purification technology developed by MicroGEM, New Zealand, with the MinION and MinIT mobile sequencing devices developed by Oxford Nanopore Technologies to produce an effective point-of-need field diagnostic system. The PDQeX extracts DNA using a cocktail of thermophilic proteinases and cell wall degrading enzymes, thermo-responsive extractor cartridges and a temperature control unit. This single-step closed system delivers purified DNA with no cross contamination. The MinIT is a newly released data processing unit that converts MinION raw signal output into base called data locally in real time, removing the need for high specification computers and large file transfers from the field. All three devices are battery powered with an exceptionally small footprint that facilitates transport and set up.

To evaluate and validate capability of the system for unbiased pathogen identification by realtime sequencing in a farmer’s field setting, we analysed samples collected from cassava plants grown by subsistence farmers in three sub-Sahara African countries (Tanzania, Uganda and Kenya). A range of viral pathogens, all with similar symptoms, greatly reduce yield or completely destroy cassava crops. 800 million people worldwide depend on cassava for food and yearly income, and viral diseases are a significant constraint on its production (https://cassavavirusactionproject.com). Early pathogen detection at a molecular level has great potential to rescue crops within a single growing season by providing results that inform decisions on disease management, use of appropriate virus resistant or replacement planting.

This case study presented conditions of working in-field with limited or no access to mains power, laboratory infrastructure, internet connectivity and highly variable ambient temperature. An additional challenge is that, generally, plant material contains inhibitors of downstream molecular processes making effective DNA purification critical. We successfully undertook real-time on-farm genome sequencing of samples collected from cassava plants on three farms, one in each country. Cassava mosaic begomoviruses were detected by sequencing leaf, stem, tuber and insect samples. The entire process, from arrival on farm to diagnosis including sample collection, processing and provisional sequencing results was complete in under 4 hours. The need for accurate, rapid and on-site diagnosis grows as globalized human activity accelerates. This technical breakthrough has applications that are relevant to human and animal health, environmental management and conservation.

## Introduction

Crop losses due to viral diseases and pests are major constraints on food security and income for millions of households in sub-Saharan Africa (SSA). Such losses can be reduced if plant diseases and pests are correctly diagnosed and identified early. To date, researchers have utilized conventional methods for definitive identification of plant viruses and their vectors in SSA including PCR, qPCR, Next Generation and Sanger sequencing, but these require laboratory infrastructure, are costly and time consuming, and can delay time-sensitive corrective actions that could be taken. Direct rapid DNA/RNA sequencing of infected material on-the-spot or near sample collection sites turns this conventional paradigm on its head by taking the laboratory closer to farmers’ fields. This reduces overall costs and gives crop protection officers and farmers in rural communities’ information that is critical for sustainable crop production and management of pests and diseases, ensuring food and income security for millions of Africans. Currently, provision of data on viruses which is essential for developing virus resistant varieties, sharing virus-indexed germplasm between regions and deployment of virus-free certified planting materials is hampered by the long time taken to receive results generated using the aforementioned conventional diagnostic methods. Our innovation will simplify information flow and fast track the deployments of virus resistant or tolerant cassava varieties directly to the farmers field. The emergence of new tools for real-time diagnostics, such as the Oxford Nanopore MinION, has proved useful for the early detection of Ebola (Quick et al. 2016) and Zika viruses (Faria et al. 2016, Quick et al. 2017). MinION consensus sequence accuracy of 99% is sufficient to identify pathogen and strain type (Calus et al. 2018). However, it can take months before results generated using other high throughput sequencing approaches (e.g. Illumina, PacBio) are available, particularly when local scientists are reliant on third-party service providers, who are often located in other countries. The delay in detecting or identifying viruses impedes quick in-situ decision making necessary for early action, crop protection advice and disease management strategies by farmers. This ultimately compounds the magnitude of crop losses and food shortages suffered by farmers. We have decreased the time to precisely detect and identify pathogens, vectors or pests, and increased resolution and reliability of results by utilizing the power of low-cost portable DNA extraction, sequencing and data analysis devices, coupled with our innovative data analysis pipelines. This real-time diagnosis in the field or located in regional laboratories quickly provides high quality and reliable diagnostics data to help farmers, seed certification agencies, scientists, crop protection and extension officers make timely and informed decisions. The immediate data accessibility makes possible dissemination of results downstream to extension officers and farmers for early disease control action via Information and Communication Technologies (ICT) applications. The application of cutting-edge sequencing technology, genomics and bioinformatics for pest and disease control has great potential to improve food security and agricultural development at large.

We propose using this technology to rapidly diagnose plant viruses and pests affecting smallholder farmers’ crops in SSA. Our case study has identified cassava DNA viruses on the farm allowing farmers, researchers and development actors to take early, positive corrective action based on rapid diagnosis of plants. This proof-of-concept shows that portable DNA sequencer technology has great potential to reduce the risk of community crop failure. We have previously conducted pilot projects in Tanzania, Uganda and Kenya testing symptomatic and asymptomatic cassava plants and already have shown that sample collection to diagnosis and results delivered to the farmer or crop protection officer can be completed within 48 hours (Boykin et al. 2018). This technology will put the power of genome sequencing directly in the hands of agriculturalists and, in the work presented here, for the first time has enabled pest and disease diagnosis within one day on-the-spot. This has significant implications for new pest and disease outbreaks, monitoring of existing disease outbreaks and biosecurity monitoring at borders between countries.

## Materials and Methods

### Tree Lab locations

Three small scale family farms growing cassava were selected, one in each of the following counties: Tanzania, Uganda and Kenya. In Tanzania (Kisamwene, Mara Region. GPS: 315N, 1°40’5”S, 33°55’55”E, 4380ft) on 1 August 2018, in Uganda (Wakiso. GPS: 255W 0°30’29”N, 32°37’19”E, 3730 ft) on 8 August 2018, and in Kenya (Kiambu. GPS: 87E 1°5’33” S, 37°19’33”E, 4570 ft) on 14 August 2018. A video of the Tanzanian Tree Lab is found here: https://vimeo.com/329068227.

### Sample selection

All samples collected are documented in Table 1. For each sequencing run we barcoded either 11, including DNA extractions from cassava leaves, stems and also whiteflies *(Bemisia tabaci)* that were found feeding on cassava mosaic disease symptomatic cassava leaves.

**Table 1.**
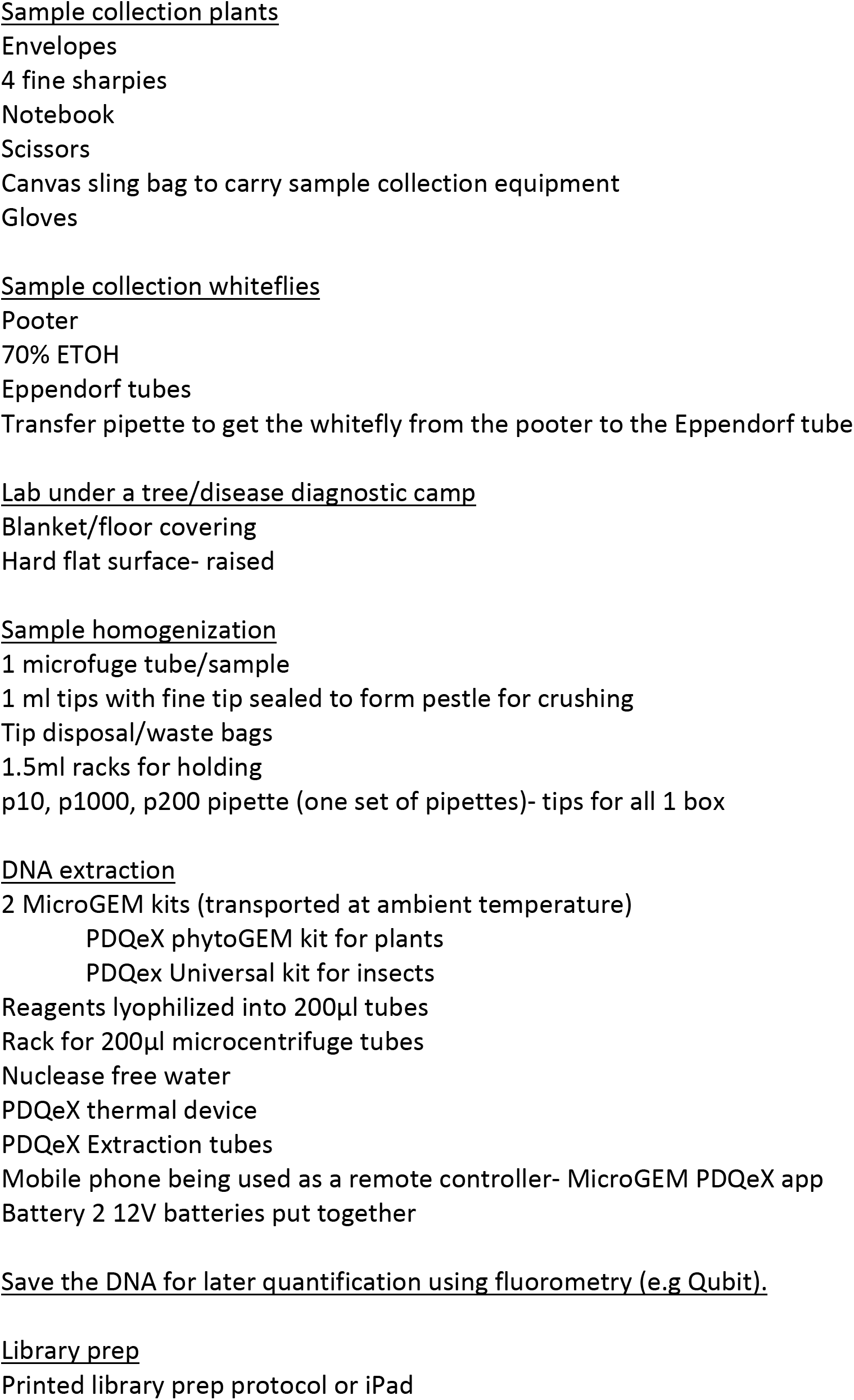
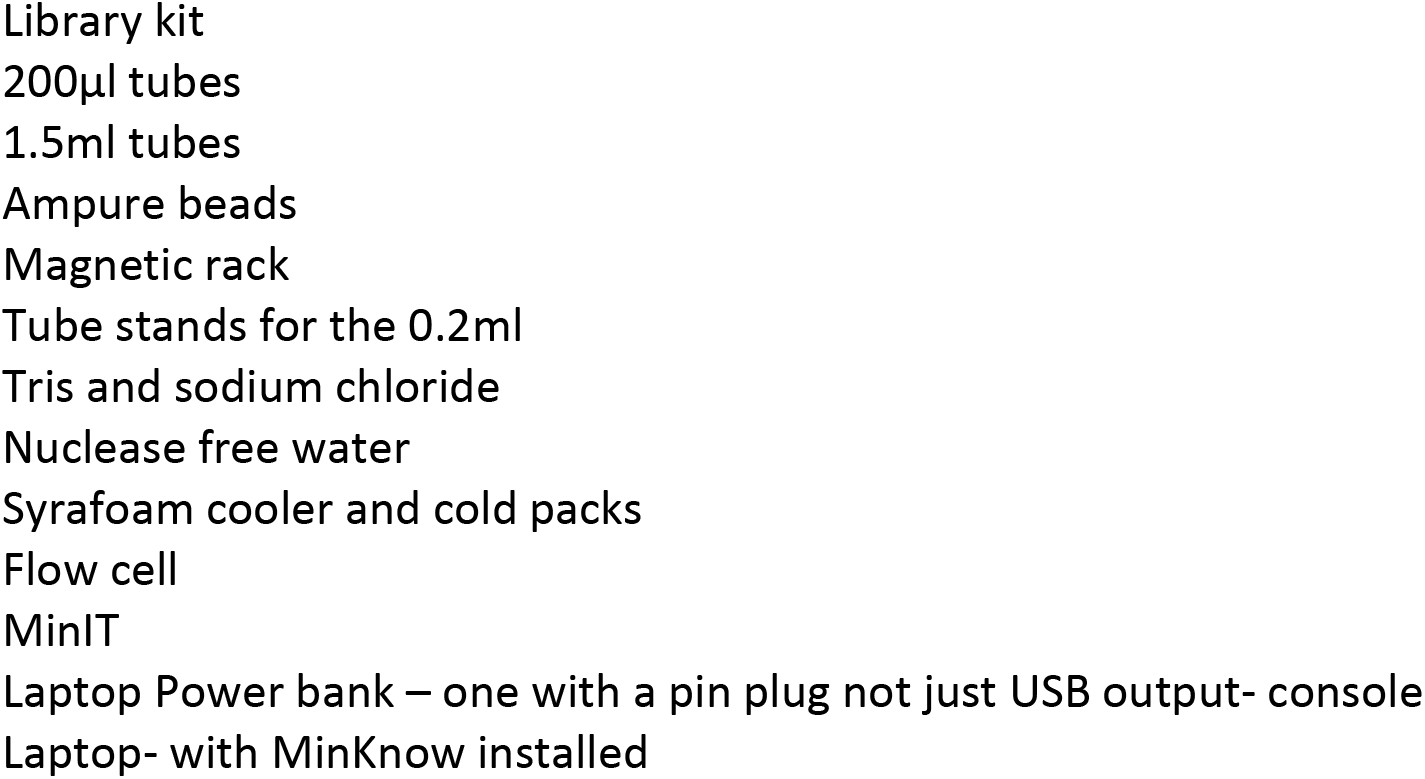
Essential equipment for Tree Lab in Tanzania, Kenya and Uganda.

### DNA extraction

The PDQeX DNA extraction system from MicroGEM NZ Ltd (Stanton et al. 2019) was used to prepare DNA from samples. Briefly, a Harris punch was used to collect four discs, 2 mm in diameter, from each leaf, stem or root sample. Homogenization was performed by hand in 1x GREEN plus buffer using a Dounce homogenizer made from sealing the end of a 1 ml pipette tip and a 1.5 ml microfuge tube. Ninety microlitres of each homogenate and 10 μl of Enhancer (MicroGEM Ltd) was added to a 200 μl tube containing a lyophilized 1x mix of the enzyme cocktail (Holmes et al. 2018); *(phytoGEM* kit, MicroGEM Ltd, New Zealand). The reaction was re-suspended by gently flicking the 200 μl tube until all reagents were well mixed. All of the reaction mix was transferred to a PDQeX extraction cartridge (Stanton et al. 2019) which was placed into the PDQeX1600 thermal incubation unit. PDQeX extraction was performed by a series of heating steps. First, incubation at 52°C for five minutes to promote cell lysis by activating cell wall degrading enzymes. Second, incubation at 75°C for five minutes to activate thermophilic proteinases to degrade sample proteins and enzymes from the previous step. Finally, heating to 95°C for 2 minutes to shrink the thermal responsive inner layer of the PDQeX extraction cartridge forcing the digested sample through a burst valve and a cleanup column into a collection tube (Stanton et al. 2019).

DNA was also extracted from whitefly. A single insect was fished from a pool of whiteflies in ethanol collected from leaves using a Pooter. The whitefly was transferred by pipette, taking as little ethanol as possible, to 98 μl of 1x BLUE buffer (MicroGEM Ltd) and pipetted up and down several times. The whole mix was added to 1x lyophilized enzyme cocktail in a 200 μl tube (prepGEM, MicroGEM Ltd). Reagents were re-suspended by gentle flicking and the contents transferred to a PDQeX extraction cartridge. The cartridge was placed in the PDQeX1600 thermal unit and heated as follows: 35°C for five minutes; 52°C for five minutes; 75°C for five minutes; 95°C for 2 minutes. DNA extraction took approximately 20 minutes in total and 7.5 μl of the collected elute was used directly for Rapid DNA library construction for MinION Sequencing. The PDQeX1600 thermal unit was powered by a 12-volt Lithium Polymer battery. The PDQeX1600 was operated using a purpose-made App from a Smart Phone that permitted run initiation, temperature profile selection and editing, and monitoring of run progress.

### Library preparation and sequencing

We utilized the Rapid Barcoding kit SQK-RBK004 with 9.4.1 flow cells (Oxford Nanopore Technologies). The SQK-RBK004 protocols were performed as described by the manufacturer (RBK_9054_v2_revB_23Jan2018). We completed the optional clean-up steps using AMPure XP beads. The 30°C and 80°C steps were performed using the PDQeX1600 thermal incubation unit. All libraries were loaded directly onto the MinION that was connected to a MinIT and live base calling was enabled. For each Tree Lab the MinION and the MinIT were plugged into a 20000mAh laptop powerbank (Comsol) set at 20V (Figure 1). The key to using a power bank for this purpose is to make sure it not only has USB inputs but also has a DC port. It ran on average 4.5 hours set on 16.5V. When the run stopped, we immediately plugged the devices into a second power bank and data generation continued.

**Figure 1.**
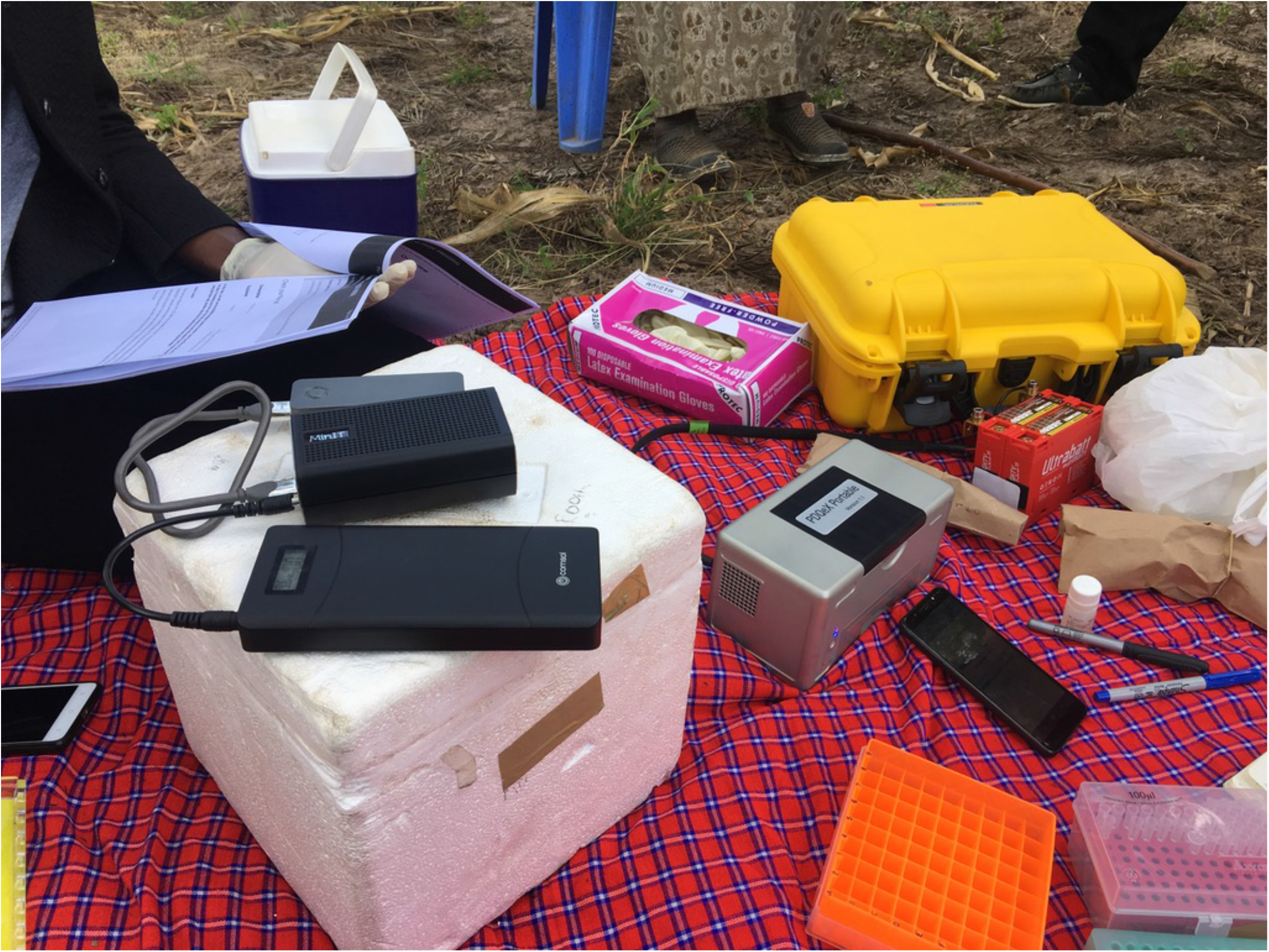
Tree Lab in Kenya. Essential equipment is listed in Table 1.

### Tree Lab data Analyses

As the data was being basecalled on the MinIT we made a test folder on the laptop called “treelab” and inside that folder we added a demultiplex folder, into which we then transferred the first two.fastq files from the MinIT into. Demultiplexing was run with Porechop (Wick 2019), preinstalled on the laptop, using the following commands >porechop-i /Users/lboykin/Desktop/treelab-b/Users/lboykin/Desktop/treelab/demultiplex. A cassava mosaic disease (CMD) reference data set was pre-curated and configured to work as a local database within Geneious vR11.1.2 (Kearse et al. 2012) (https://figshare.com/articles/Nanopore_sequencing_of_cassava_from_Tanzania_Uganda_and_Kenya/6667409). Twelve folders were created in Geneious, and the associated .fastq files from the “demultiplex” folder were drag and dropped into the relevant folder created within Geneious. BLASTn (Altschul et al. 1990) analysis was performed, ensuring the local CMD database was specified. The results from the search against the CMD database were visualized in-situ within Geneious and discussed with farmers and extension workers.

### Post Tree Lab data analyses

Scripts from David Eccles’ [Bioinformatics Scripts repository] (https://doi.org/10.5281/zenodo.596663) were used to carry out subsequent read QC and analysis. Sequenced read lengths were measured using [fastx-length.pl], and these lengths were used to generate length-based QC plots using [length_plot.r].

### Assembly

To determine whether any barcoded read sets could be assembled, an initial assembly attempt was made on each subset using Canu v1.8, with a genome size of 400M, ignoring any warning messages about coverage being too low. The large genome size ensured that no reads are discarded and suppressing the coverage warning ensured that Canu would attempt an assembly with all the available reads. Previous discussions with Canu developer Sergey Koren (pers. comm.) indicated that adjusting the target genome size had no other effect on the contig assemblies that Canu produces.

### Blast

We confirmed these in-field results by performing a post diagnostic blast of reads on the Nimbus Cloud at the Pawsey Supercomputing Center with blast 2.2.31 against the full NCBI nucleotide database to confirm results. For reference the specific database used was {$ blastcmd –db nt/nt –info} Database: Nucleotide collection 49,266,009 sequences; 188,943,333,900 total bases Date: Aug 8, 2018 12:38 PM. The data were processed into a blast archive using a blast script with the following parameters (Script attached) {$blastn-query “$file” -db /mnt/nucdb/nt/nt -outfmt 11 -culling_limit 10 -out “out.$file.asn” -num_threads 17 } then converted into XML (for loading into Geneious) and HTML for viewing.

### Blastn analysis – MEGAN

Blastn results produced from the Nimbus cloud analysis pipeline were also visualized using MEGAN Community Edition version 6.12.6 (Huson et al. 2016) on the Zeus computing resource located at the Pawsey Supercomputing Center.

### Blastn analysis – Kraken

We used kraken2 [https://github.com/DerrickWood/kraken2] to classify demultiplexed reads using the Loman Lab “maxikraken2” database [https://lomanlab.github.io/mockcommunity/mc_databases.html#maxikraken2_1903_140gb-march-2019-140gb], on the Zeus computing resource located at the Pawsey Supercomputing Center.

## Results

### Tree Lab

All essential equipment that were used in the Tree Labs in Kenya, Uganda and Tanzania are listed in Table 1 and shown in Figure 1. Summary statistics for our three Tree Labs are shown in Table 2 and Figure 2. Table 2 summarizes DNA sequencing metrics from all three Tree Lab experiments. Each MinION run contained 11 barcoded libraries representing 11 individual samples. All DNA samples, except the ACMV and EACMV controls (Tanzania) were extracted using the PDQeX system with 11 samples prepared in lab from exemplar material collected from scientific plots and 20 DNA samples extracted on farm. A total of 1,442,599 sequences were produced across all the experiments. Of these, barcodes could only be resolved for 550,938 sequences using Porechop to demultiplex the samples. Mean sequence length across all sequencing runs ranged from 355bp to 948bp with the longest read being 276,793 bp.

**Figure 2.**
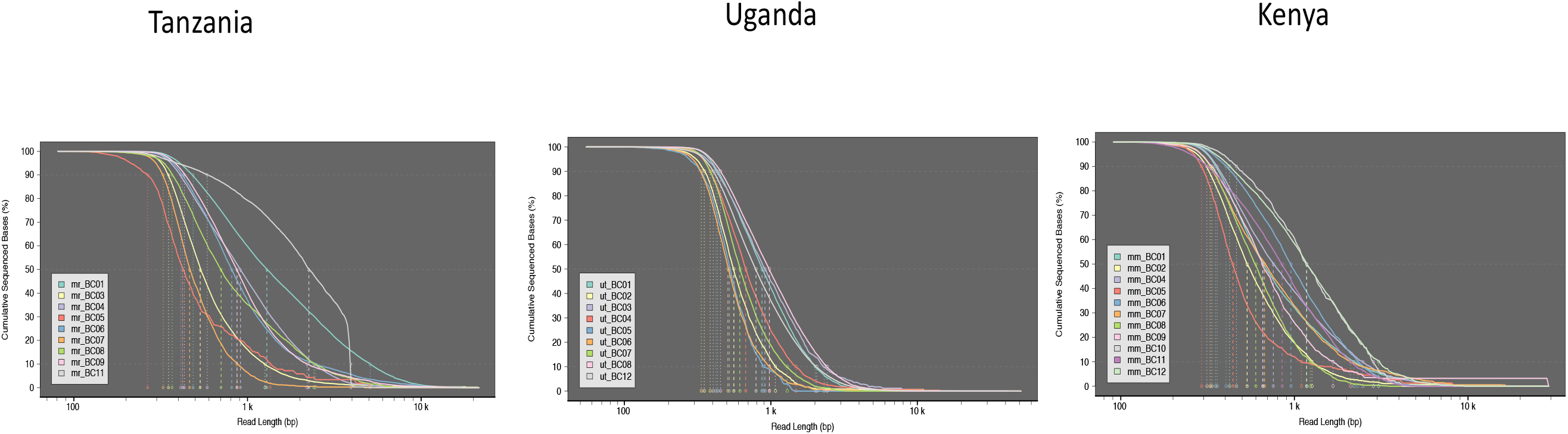
Cumulative density curves showing the proportion of sequenced bases with length greater than a particular length (with L10/L50/L90 highlighted).

**Table 2.** Summary statistics and locality information for the three Tree Labs in Tanzania, Kenya and Uganda. * indicates DNA extraction carried out using PDQeX in the laboratory before sequencing under the tree.

| Sample | Barcode | Variety | Severity Score | Tissue type | Total reads | Max. Seq Length | CMV Blast hits | Max. CMV length | Min CMV length | % CMV reads | Canu Contigs |
| --- | --- | --- | --- | --- | --- | --- | --- | --- | --- | --- | --- |
| **Tanzania** |  |  |  |  |  |  |  |  |  |  |  |
| 1 | 1 | Kilati - Local Variety | 4 | Leaf | 61,317.0 | 26,107.0 | 125.0 | 2,808.0 | 196.0 | 0.204 | 0 |
| 3 | 3 | Kilati - Local Variety | 4 | Leaf | 21,468.0 | 16,098.0 | 14.0 | 815.0 | 288.0 | 0.065 | 0 |
| 4 | 4 | Kasuxsali - Local Variety | 3 | Leaf | 31,300.0 | 17,624.0 | 21.0 | 941.0 | 129.0 | 0.067 | 12 |
| 5 | 5 | Mukombozi - Virus Resistant | 1 | Leaf | 2,117.0 | 24,665.0 | 7.0 | 815.0 | 198.0 | 0.331 | 0 |
| 6 | 6 | Mukombozi - Virus Resistant | 1 | Leaf | 27,178.0 | 12,340.0 | 8.0 | 826.0 | 85.0 | 0.029 | 0 |
| 7 | 7 | Mukombozi - Virus Resistant | 1 | Leaf | 5,634.0 | 17,003.0 | 5.0 | 484.0 | 243.0 | 0.089 | 0 |
| 8 | 8 | Whitefly close to Mukombozi - Virus Resistant | 1 | 1x Whitefly, in EtOH | 6,237.0 | 8,753.0 | 26.0 | 828.0 | 52.0 | 0.417 | 0 |
| 9 | 9 | Whitefly close to Mukombozi - Virus Resistant | 1 | 1x Whitefly, in EtOH | 25,289.0 | 17,900.0 | 3.0 | 0.0 | 0.0 | 0 | 2 |
| 10 | 10 | Whitefly close to Mukombozi - Virus Resistant | 1 | 1x Whitefly, in EtOH | 798.0 | 23,259.0 | 21.0 | 815.0 | 187.0 | 2.632 | 0 |
| 11* | 11 | ACMV - Positive Control DNA |  |  | 10,966.0 | 28,541.0 | 4,311.0 | 1,598.0 | 31.0 | 39.312 | 0 |
| 12* | 12 | EACMV - Positive Control DNA |  |  | 1,797.0 | 22,871.0 | 9.0 | 830.0 | 191.0 | 0.501 | 0 |
| None | None | Porechop unable to match to Rapid Barcode |  |  | 449,981.0 | 276,793.0 | 8,136.0 | 2,757.0 | 28.0 | 1.808 |  |
| **Uganda** |  |  |  |  |  |  |  |  |  |  |  |
| 1* | 1 | R39-B1-UG15F289P503 | 4 | Leaf from stem 1 | 24,343.0 | 24,373.0 | 20.0 | 1,222.0 | 123.0 | 0.082 | 2 |
| 1.1* | 2 | R39-B1-UG15F289P503 | 4 | Phloem, stem 1 - top | 5,135.0 | 50,184.0 | 0.0 | 0.0 | 0.0 | 0 | 0 |
| 1.2* | 3 | R39-B1-UG15F289P503 | 4 | Phloem, stem 1 - mid | 4,010.0 | 9,768.0 | 0.0 | 0.0 | 0.0 | 0 | 0 |
| 1.3* | 4 | R39-B1-UG15F289P503 | 4 | Phloem, stem 1 - bottom | 10,314.0 | 48,265.0 | 1.0 | 402.0 | 402.0 | 0.010 | 0 |
| 5 | 5 | Kwatempale from Sarah's Farm | 4 | Leaf | 3,012.0 | 66,062.0 | 11.0 | 604.0 | 107.0 | 0.365 | 0 |
| 6 | 6 | Kwatempale from Sarah's Farm | 5 | Leaf | 5,074.0 | 5,243.0 | 1.0 | 235.0 | 235.0 | 0.020 | 0 |
| 7 | 7 | Wild Plant from Naomi's Farm | 3 | Leaf | 16,386.0 | 45,024.0 | 6.0 | 1,121.0 | 352.0 | 0.037 | 0 |
| 9 | 8 | Sick branch from NAROCass1 from Naomi's Farm | 3 | Leaf | 37,822.0 | 20,509.0 | 15.0 | 2,021.0 | 166.0 | 0.040 | 12 |
| WF5 | 9 | Whitefly from sample 5 | 4 | 1x Whitefly, no EtOH | 347.0 | 41,599.0 | 0.0 | 0.0 | 0.0 | 0 | 0 |
| WF7 | 10 | Whitefly from sample 7 | 3 | 1x Whitefly, no EtOH | 2,613.0 | 2,613.0 | 0.0 | 0.0 | 0.0 | 0 | 0 |
| WF8 | 11 | Whitefly from sample 8 | 1 | 1x Whitefly, no EtOH | 705.0 | 15,596.0 | 0.0 | 0.0 | 0.0 | 0 | 0 |
| None* | None | Porechop unable to match to Rapid Barcode |  |  | 196,290.0 | 267,436.0 | 59.0 | 1,677.0 | 42.0 | 0.030 |  |
| **Kenya** |  |  |  |  |  |  |  |  |  |  |  |
| 1 | 1 | Local | 1 | Leaf | 43,049.0 | 39,851.0 | 1.0 | 123.0 | 123.0 | 0.002 | 1 |
| 2 | 2 | Local | 1 | Leaf | 15,890.0 | 66,540.0 | 0.0 | 0.0 | 0.0 | 0 | 0 |
| 4 | 4 | Local | 1 | Leaf | 38,291.0 | 53,843.0 | 3.0 | 890.0 | 251.0 | 0.008 | 1 |
| 5 | 5 | Local | 1 | Leaf | 9,213.0 | 44,230.0 | 0.0 | 0.0 | 0.0 | 0 | 0 |
| L1* | 6 | Stem from CMV infected plant | 4 | Leaf | 28,450.0 | 18,287.0 | 219.0 | 2,228.0 | 98.0 | 0.770 | 3 |
| L2* | 7 | Stem from CMV infected plant | 4 | Leaf | 17,320.0 | 42,191.0 | 76.0 | 2,127.0 | 78.0 | 0.439 | 0 |
| S1* | 8 | Stem from CMV infected plant | 4 | Phloem, 22.5cm from tip | 16,310.0 | 24,566.0 | 10.0 | 1,485.0 | 269.0 | 0.061 | 0 |
| S2* | 9 | Stem from CMV infected plant | 4 | Phloem, 52.3cm from tip | 5,346.0 | 54,245.0 | 0.0 | 0.0 | 0.0 | 0 | 0 |
| R1* | 10 | Stem from CMV infected plant | 4 | Root 1, under outer bark | 7,336.0 | 15,689.0 | 0.0 | 0.0 | 0.0 | 0 | 0 |
| R2* | 11 | Stem from CMV infected plant | 4 | Root 2, under outer bark | 21,576.0 | 24,853.0 | 0.0 | 0.0 | 0.0 | 0 | 0 |
| H1* | 12 | Leaf from Healthy Plant | 1 | Leaf | 44,295.0 | 33,307.0 | 0.0 | 0.0 | 0.0 | 0 | 3 |
| None | None | Porechop unable to match to Rapid Barcode |  |  | 245,390.0 | 265,898.0 | 75.0 | 2,404.0 | 28.0 | 0.031 |  |
| **TOTALS** |  |  |  |  | **492,466.0** | **56,958.3** |  |  |  |  |  |

Raw reads of Cassava mosaic begomoviruses (CMBs) sequences were detected in 21 samples with the longest CMB read reaching 2808 bp, close to the full genome size. A total of 18 leaf samples were sequenced of which 15 were found to contain CMBs. Two of the 5 stem samples sequenced were found to contain CMBs whereas neither of the two root samples sequenced presented CMB sequences. Six single whiteflies were tested with 2 being positive for CMBs. All libraries, regardless of CMB content produced DNA reads indicating that sequencing was successful for all samples. CMBs were detected in plants with symptoms and there was a suggestion that the number of CMB sequences detected possibly correlated with symptom severity scores, but more data will be required to prove this. Interestingly, a known healthy plant taken from the scientific plot at JKUAT did not yield CMB sequences (Table 2). Following assembly with Canu 8 of the 21 samples gave complete assembled virus genomes, however, gave less than 10 fold coverage.

### Post Tree Lab data analyses

We investigated whether there was any effect of sample type on read length. The cumulative density curves (Figure 2) show the proportion of sequenced bases with length greater than a particular length (with L10/L50/L90 highlighted). Additional length-based QC plots can be found in the supplemental information (Supplemental File 1).

### MEGAN results

The primary targets of this analysis were known cassava viruses, as well as the host, either cassava plant *(Manihot esculenta)* or the whitefly *(Bemisia tabaci)* and its endosymbionts. Results are summarized in Table 3, and in general the desired result of virus (EACMV or ACMV) and host DNA were recovered from all symptomatic samples.

**Table 3.** Megan Results of Blastn from Nimbus cloud.

|  | Total reads | Reads classified | Manihot esculenta | EACMV | ACMV | TLCV | Begomo-associated DNA-III | Bemisia tabaci | Bemisia afer | Candidatus Portiera aleyrodidarum | Other |
| --- | --- | --- | --- | --- | --- | --- | --- | --- | --- | --- | --- |
| **Tanzania** |  |  |  |  |  |  |  |  |  |  |  |
| 1 | 61,317 | 23,481 | 15,862 | 63 | - | 1 |  |  |  |  |  |
| 3 | 21,468 | 5,407 | 3,509 | 5 |  |  |  |  |  |  |  |
| 4 | 31,309 | 10,561 | 6,462 | 2 | 5 | 1 |  |  |  |  |  |
| 5 | 2,117 | 325 | 144 |  | 2 |  |  | 3 |  | 3 |  |
| 6 | 27,178 | 2,127 | 76 |  | 1 |  |  | 521 | 67 | 802 |  |
| 7 | 5,634 | 1,141 | 669 |  | 1 |  |  |  |  |  |  |
| 8 | 6,237 | 1,449 | 88 |  | 5 |  |  | 912 |  | 15 |  |
| 9 | 25,289 | 2,303 | 126 | 2 |  |  |  | 506 | 57 | 905 |  |
| 10 | 789 | 166 | 66 |  | 3 |  |  | 11 |  | 2 |  |
| 11 | 10,966 | 7,843 | 69 | 3 | 616 |  |  |  |  | 2 |  |
| 12 | 1,797 | 356 | 171 |  | 2 |  |  | 13 |  | 7 |  |
| **Uganda** |  |  |  |  |  |  |  |  |  |  |  |
| 1 | 18,853 | 5,662 | 3,073 |  | 11 |  | 1 |  |  |  |  |
| 1.1 | 5,135 | 876 | 402 |  |  |  |  |  |  |  |  |
| 1.2 | 4,010 | 1,034 | 591 |  |  |  |  |  |  |  |  |
| 1.3 | 10,314 | 1,933 | 864 |  | 1 |  |  |  |  |  |  |
| 5 | 3,012 | 556 | 255 | 1 | 7 |  |  |  |  |  |  |
| 6 | 5,074 | 758 | 39 |  | 1 |  |  |  |  |  |  |
| 7 | 16,386 | 4,608 | 2,666 |  | 4 |  |  |  |  |  |  |
| 9 | 37,822 | 12,768 | 7,268 | 10 |  |  |  |  |  |  |  |
| WF5 | 347 | 51 | 20 |  |  |  |  | 10 |  |  |  |
| WF7 | 243 | 48 | 21 |  |  |  |  |  |  |  |  |
| WF8 | 705 | 128 | 35 |  |  |  |  | 9 |  | 1 |  |
| **Kenya** |  |  |  |  |  |  |  |  |  |  |  |
| 1 | 43,049 | 10,283 | 9,947 |  | 1 | 1 | 1 |  |  |  |  |
| 2 | 15,890 | 2,968 | 2,854 |  |  |  |  |  |  |  |  |
| 4 | 38,291 | 8,648 | 8,836 | 1 | 1 |  |  |  |  |  |  |
| 5 | 8,213 | 1,959 | 1,887 |  |  |  |  |  |  |  |  |
| L1 | 28,450 | 9,968 | 9,580 | 45 | 81 |  |  |  |  |  | SLCV (1) |
| L2 | 17,320 | 4,367 | 2,050 | 29 | 4 |  |  |  |  |  |  |
| S1 | 16,310 | 3,718 | 1,933 | 5 |  | 1 | 1 |  |  |  |  |
| R1 | 7,336 | 2,303 | 1,016 |  |  |  |  |  |  |  |  |
| R2 | 21,578 | 6,810 | 2,000 |  |  |  |  |  |  |  |  |
| H1 | 44,295 | 15,537 | 8,389 |  |  |  | 1 |  |  |  |  |

### Kraken2 results

The analysis using Kraken2 had an approximately 50% classification success rate (IQR 45-52% unclassified reads). This database is for human + microbial and viral sequence, so any eukaryote reads (e.g. from cassava or whitefly) would probably be unclassified by Kraken2 or assigned to the human taxa. The sample with the highest classification success was the ACMV positive control from Tanzania (mr_BC11, 5.6% unclassified), and the lowest classification success was the leaf tissue sample #2 from Kenya (mm_BC02, 61% unclassified).

Results were aggregated into a table using Pavian (Supplemental File 2) to identify common elements of each sample. Begomovirus reads were detected in 15 samples, with very high proportions of Begomovirus (8.6%) in the ACMV Positive control from Tanzania (mr_BC11), and above-average proportions (0.25%) in Kwatempale sample #5 from Sarah’s Farm in Uganda (ut_BC05). ACMV and EACMV were detected in 11 samples.

## Discussion

This case study was designed to show the possibility to go from sample to diagnosis, in a regional setting, on farm in three hours versus the normal 6 months with conventional methods. The results of this research show that it is indeed possible, and that it is possible to use a range of battery powered devices to achieve DNA extraction, long read sequencing and analysis all under a tree on the farm while the farmers wait for results.

Access to next generation sequencing technology, or to services that offer access has been a major barrier to their use in diagnostics for scientists, and particularly many agricultural scientists in SSA. The advent of the Oxford Nanopore Technologies MinION has brought this technology to their door in recent years, and with access to training through various institutions and especially the Oxford Nanopore Technologies run “Pore Safari” there are more and more users in the region. Previous studies that have used the technology for real time analysis of pathogen outbreaks, such as the Ebola and Zika studies (Faria et al. 2016, Quick et al. 2016, Quick et al. 2017) have still relied heavily on the transport of bulky laboratory equipment, or local acquisition of it to perform their work. Previous work by our team of scientists (Boykin et al. 2018) showed that the turnaround time to result could be 48 hours, and now with the addition of the PDQeX and MinIT to the system we have been able to reduce the time to 4 hours and perform the entire process in the field and under a tree.

One of the major barriers to producing these outputs in the field has until now been the lack of a simple, quick and effective methods to extract DNA from a sample without the need for laboratory equipment requiring mains power and space, items such as benchtop centrifuges, fridges, freezers and temperature sensitive extraction kits which can be bulky and rely on traditional laboratory infrastructure. The use of the PDQeX in the system described in this case study was the real game changer: compact and able to operate from a battery, it made nucleic acid extraction possible.

This study also highlighted where the next gains for in-field sequencing are to be made, as improvements are required in rapid data analysis. The MinIT eliminated obstacles to base calling, by converting the raw reads into .fast5 and .fastq reads in real time. This moves the data analysis bottleneck in the pipeline to the Blast analysis. Blast is not fast analysis, and so for now we must rely on a pre-curated database of known or expected pathogens and host genomes. This poses risks, in that new and emerging pathogens or vectors could be missed in the first instance, and not seen until subsequent data analysis when the scientist has returned to the lab or is within range of a good internet connection capable of uploading large amounts of data to the cloud. In our case, we can predict what sorts of genomes should be in our custom database, but for use in biosecurity, and at borders between countries a better solution is required.

Read length distributions were generally quite similar for all samples. The Tanzania samples showed the greatest difference in read length distribution (L50 range 400 - 2000 bp). The ACMV Positive control from Tanzania (mr_BC11) showed a very pronounced spike of reads at around 4kb (presumably near full-length ACMV sequence). Apart from that sample, there was no obvious association between read length distribution and tissue type or variety. The Uganda samples had a moderate read length distribution spread (L50 range 500-1000 bp), which split into two clusters of slightly shorter and longer reads (BC02, BC04, BC05, BC06, BC07; BC01, BC03, BC08, BC12). These clusters did not appear to have any relationship with tissue type or variety. The Kenya samples had a similarly moderate read length distribution spread (L50 range 500-1000 bp), with no obvious clustering, or association of distribution with tissue type or variety.

MinION Rapid libraries use transposase to fix sequencing adaptors to DNA fragments. The ratio of DNA to transposase complex for the MinION Rapid kit has been optimized for 400 ng DNA and at lower amounts DNA is susceptible to over fragmentation. This may account for DNA fragment length falling around 900 bp, however, the control DNA also gave similar read length characteristics. Though there was not enough data collected on farm for a thorough statistical analysis these results did show both yield and integrity of the DNA extracted using the PDQeX was of sufficient quality for diagnostic sequencing. We successfully retrieved enough data from each sample to establish whether the virus in the plant was EACMV or ACMV. Assembly with Canu suggested that in this case, while there was enough data to assemble whole genomes, the average coverage meant the quality was not sufficient for downstream applications such as recombination detection and other evolutionary analyses. We anticipate that as the DNA extraction methods improve, and in field library preparation becomes easier this will be possible. An alternative would be to investigate the use of a panel-like targeted amplicon approach or CRISPR/Cas9 enrichment, but again this removes the likelihood of detecting unknowns in the samples, and could lead to samples giving negative results not being followed up, or the time to result being blown out to days or weeks if they need to return to a laboratory to complete a different type of library preparation.

Compared with other in-field diagnostic tools, this system involving the MinION is unique in its ability to detect anything that might be present in the sample. Other in-field diagnostic tools, including serological based dipsticks, LAMP-PCR, in field qPCR and AI driven applications on smart phones all have one single thing in common – they require a prior knowledge of the suspected pathogen, coupled with targeted design of antibodies, primers or training for known positives to function effectively. The only decision required to run the MinION is whether to prepare a DNA or an RNA library.

## Executive summary

**Can we go from sample to answer on the farm?** Yes

**DNA extraction to library prep to sequencing?** Yes

**Can we detect virus in leaves off the grid at the farm?** Yes

**Can we detect virus in whiteflies off the grid on the farm?** Yes

**Can we detect virus in stems off the grid on the farm?** Yes

**Do we get enough to coverage of the viral genomes to generate polished genomes to track the evolution of the viruses real-time?** No

**Video of Tree Lab:** https://vimeo.com/329068227

## Acknowledgements

We are grateful to the farmers in Tanzania, Kenya and Uganda who allowed us access to their farms. This work is dedicated to them. We were honored to have worked along side scientist Jimmy Akono who passed earlier this year in Uganda. Special thank you to our drivers Honest Kway, Benson Ongori, and Jimmy Sebayiga. Filming was done by Andrew Court. PDQeX was developed under MBIE UOOX1507, New Zealand. Resources provided by Pawsey Supercomputing Centre with Funding from the Australia Government and the Government of Western Australia supported computation analysis.

## References

Altschul, S. F., W. Gish, W. Miller, E. W. Myers, and D. J. Lipman. 1990. Basic local alignment search tool. J Mol Biol. 215:403–410.

Boykin, L. M., A. Ghalab, B. R. De Marchi, A. Savill, J. M. Wainaina, T. Kinene, S. Lamb, M. Rodrigues, M. Kehoe, J. Ndunguru, F. Tairo, P. Sseruwagi, C. Kayuki, D. Mark, J. Erasto, H. Bachwenkizi, T. Alicai, G. Okao-Okuja, P. Abidrabo, E. Ogwok, J. F. Osingada, J. Akono, E. Ateka, B. Muga, and Kiarie. 2018. Real time portable genome sequencing for global food security. F1000 7:1101.

Calus, S. T., U. Z. Ijaz, and A. J. Pinto. 2018. NanoAmpli-Seq: a workflow for amplicon sequencing for mixed microbial communities on the nanopore sequencing platform. Gigascience 7.

Faria, N. R., E. C. Sabino, M. R. Nunes, L. C. Alcantara, N. J. Loman, and O. G. Pybus. 2016. Mobile realtime surveillance of Zika virus in Brazil. Genome Med 8:97.

Holmes, A. S., M. G. Roman, and S. Hughes-Stamm. 2018. In-field collection and preservation of decomposing human tissues to facilitate rapid purification of STR typing. Forensic Sci. Int. Genet. 36:124–129.

Huson, D. H., S. Beier, I. Flade, A. Gorska, M. El-Hadidi, S. Mitra, H. J. Ruscheweyh, and R. Tappu. 2016. MEGAN Community Edition-Interactive Exploration and Analysis of Large-Scale Microbiome Sequencing Data. PLoS Comput Biol 12:e1004957.

Quick, J., N. D. Grubaugh, S. T. Pullan, I. M. Claro, A. D. Smith, K. Gangavarapu, G. Oliveira, R. Robles-Sikisaka, T. F. Rogers, N. A. Beutler, D. R. Burton, L. L. Lewis-Ximenez, J. G. de Jesus, M. Giovanetti, S. C. Hill, A. Black, T. Bedford, M. W. Carroll, M. Nunes, L. C. Alcantara, Jr., E. C. Sabino, S. A. Baylis, N. R. Faria, M. Loose, J. T. Simpson, O. G. Pybus, K. G. Andersen, and N. J. Loman. 2017. Multiplex PCR method for MinION and Illumina sequencing of Zika and other virus genomes directly from clinical samples. Nat Protoc 12:1261–1276.

Quick, J., N. J. Loman, S. Duraffour, J. T. Simpson, E. Severi, L. Cowley, J. A. Bore, R. Koundouno, G. Dudas, A. Mikhail, N. Ouedraogo, B. Afrough, A. Bah, J. H. Baum, B. Becker-Ziaja, J. P. Boettcher, M. Cabeza-Cabrerizo, A. Camino-Sanchez, L. L. Carter, J. Doerrbecker, T. Enkirch, I. G. G. Dorival, N. Hetzelt, J. Hinzmann, T. Holm, L. E. Kafetzopoulou, M. Koropogui, A. Kosgey, E. Kuisma, C. H. Logue, A. Mazzarelli, S. Meisel, M. Mertens, J. Michel, D. Ngabo, K. Nitzsche, E. Pallash, L. V. Patrono, J. Portmann, J. G. Repits, N. Y. Rickett, A. Sachse, K. Singethan, I. Vitoriano, R. L. Yemanaberhan, E. G. Zekeng, R. Trina, A. Bello, A. A. Sall, O. Faye, O. Faye, N. Magassouba, C. V. Williams, V. Amburgey, L. Winona, E. Davis, J. Gerlach, F. Washington, V. Monteil, M. Jourdain, M. Bererd, A. Camara, H. Somlare, A. Camara, M. Gerard, G. Bado, B. Baillet, D. Delaune, K. Y. Nebie, A. Diarra, Y. Savane, R. B. Pallawo, G. J. Gutierrez, N. Milhano, I. Roger, C. J. Williams, F. Yattara, K. Lewandowski, J. Taylor, P. Rachwal, D. Turner, G. Pollakis, J. A. Hiscox, D. A. Matthews, M. K. O’Shea, A. M. Johnston, D. Wilson, E. Hutley, E. Smit, A. Di Caro, R. Woelfel, K. Stoecker, E. Fleischmann, M. Gabriel, S. A. Weller, L. Koivogui, B. Diallo, S. Keita, A. Rambaut, P. Formenty, S. Gunther, and M. W. Carroll. 2016. Real-time, portable genome sequencing for Ebola surveillance. Nature 530:228–232.

Stanton, J. L., A. Muralidhar, C. J. Rand, and D. J. Saul. 2019. Rapid extraction of DNA suitable for NGS workflows from bacterial cultures using the PDQeX.. BioTechniques 66:1–7.

