## Supplemental File 1 for "Tree Lab: Portable genomics for early detection of plant viruses and pests in Sub-Saharan Africa"

Supplement File 1.

Decoding: mr=Tanzania, ut=Uganda and mm=Kenya. BC number correspond to the barcode in Table 2.

Scripts from David Eccles' [Bioinformatics Scripts repository] (https://doi.org/10.5281/zenodo.596663) were used to carry out subsequent read QC and analysis. Sequenced read lengths were measured using [fastx-length.pl], and these lengths were used to generate length-based QC plots using [length_plot.r]. The produced plots are as follows:

1. Curves showing the number of sequenced *reads* within each length bin
2. Curves showing the number of sequenced *bases* within each length bin
3. Density curves showing the proportion of sequenced bases within each length bin
4. Cumulative density curves showing the proportion of sequenced bases with length greater than a particular length (with L10/L50/L90 highlighted)
5. Digital electrophoresis plot representing graph #2
6. Digital electrophoresis plot representing graph #3

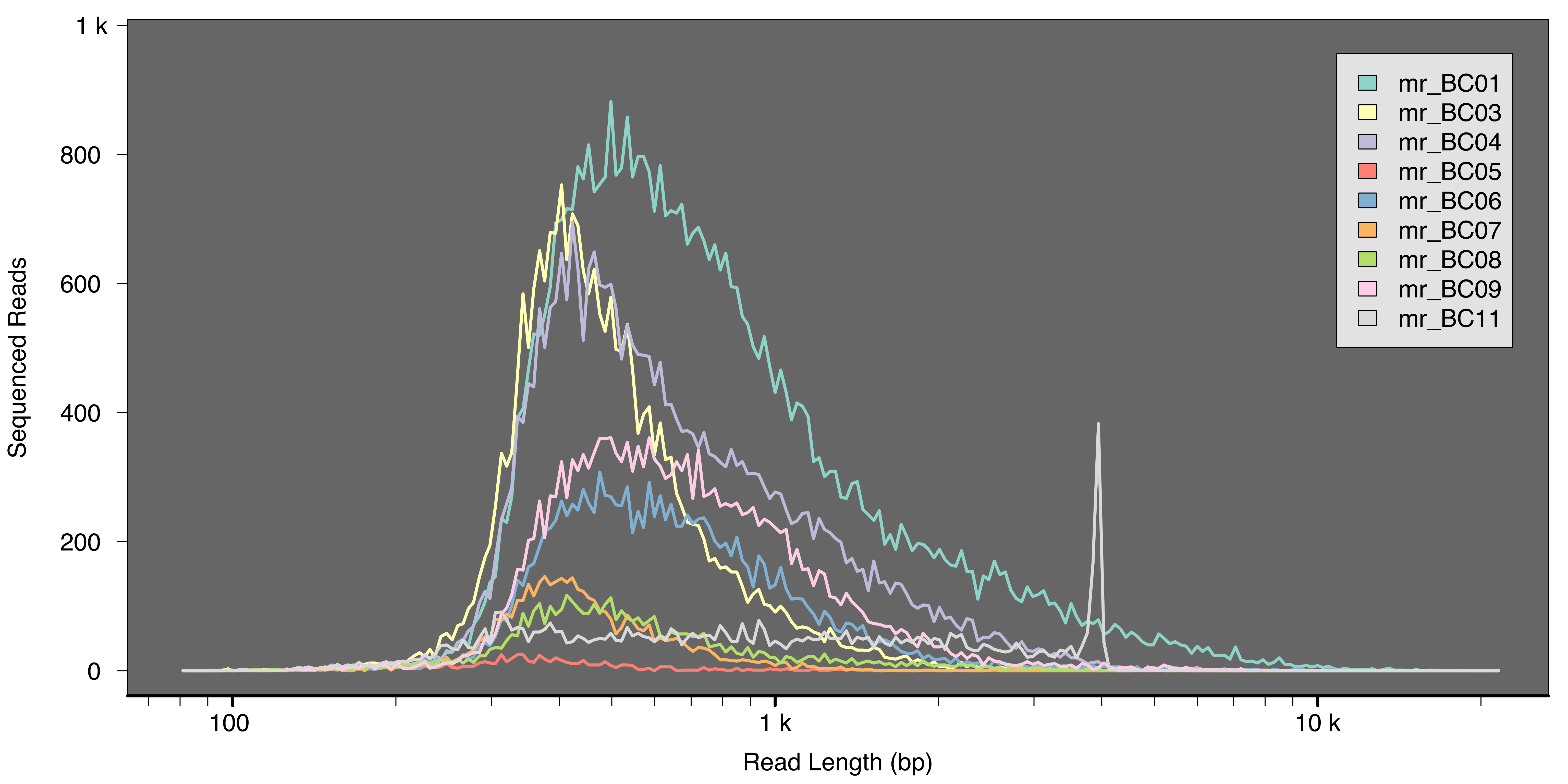
1 k
800
600
400
200
0
Sequenced Reads
100
1 k
10 k
Read Length (bp)
mr_BC01
mr_BC03
mr_BC04
mr_BC05
mr_BC06
mr_BC07
mr_BC08
mr_BC09
mr_BC11

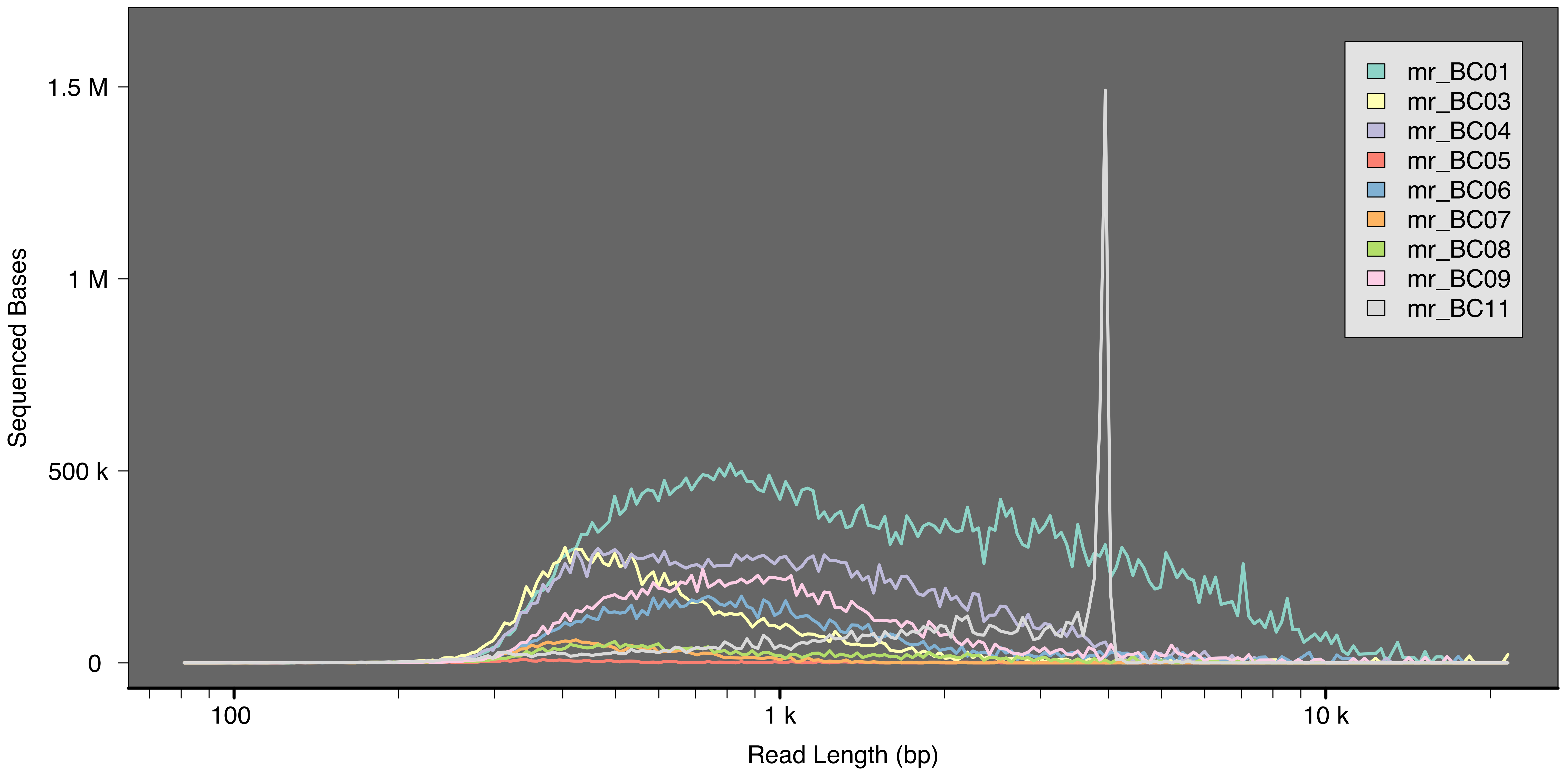
Sequenced Bases
1.5 M
1 M
500 k
0
100
1 k
10 k
Read Length (bp)
mr_BC01
mr_BC03
mr_BC04
mr_BC05
mr_BC06
mr_BC07
mr_BC08
mr_BC09
mr_BC11

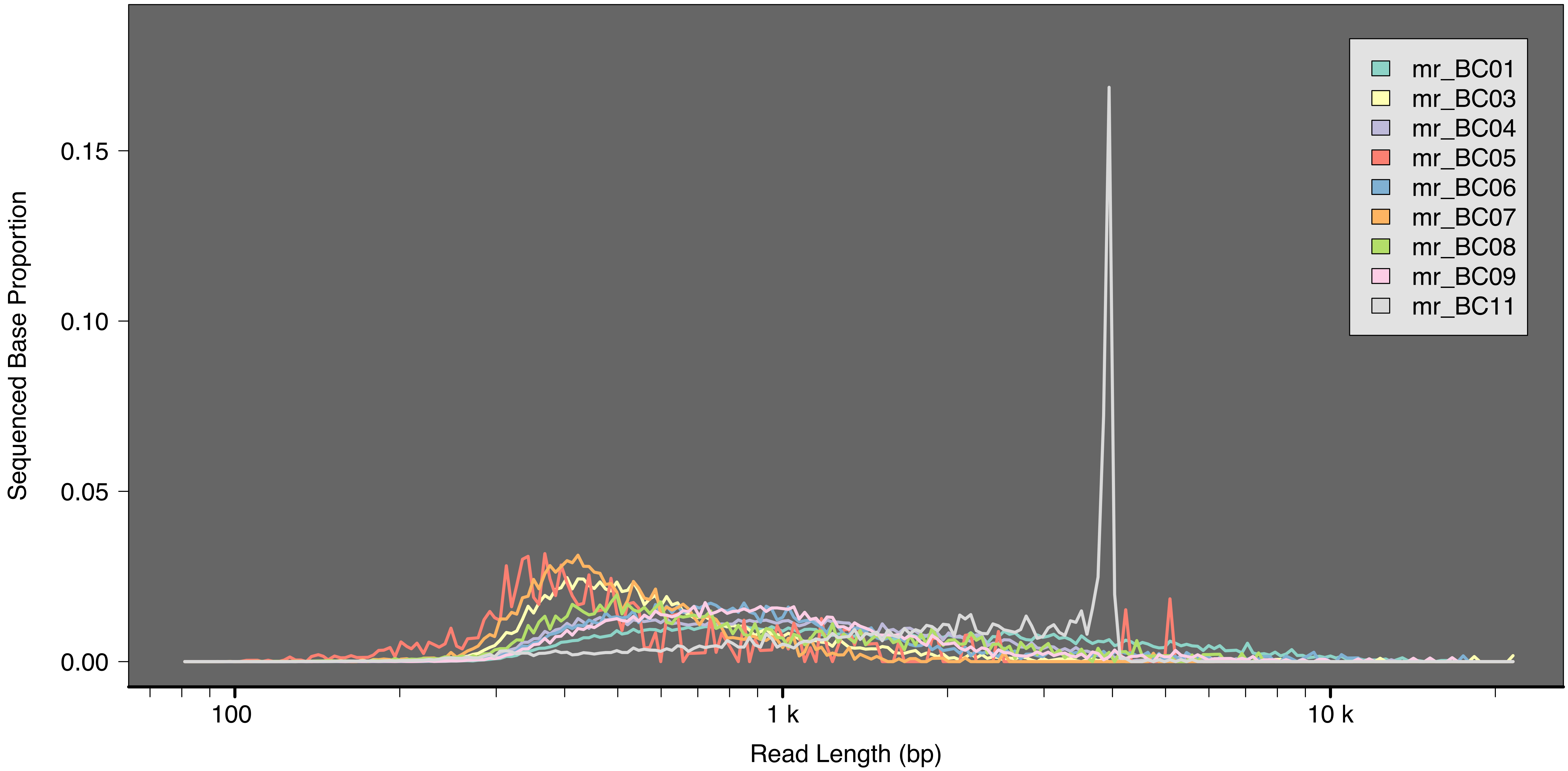

Sequenced Base Proportion
0.15
0.10
0.05
0.00
100
1 k
10 k
Read Length (bp)
mr_BC01
mr_BC03
mr_BC04
mr_BC05
mr_BC06
mr_BC07
mr_BC08
mr_BC09
mr_BC11

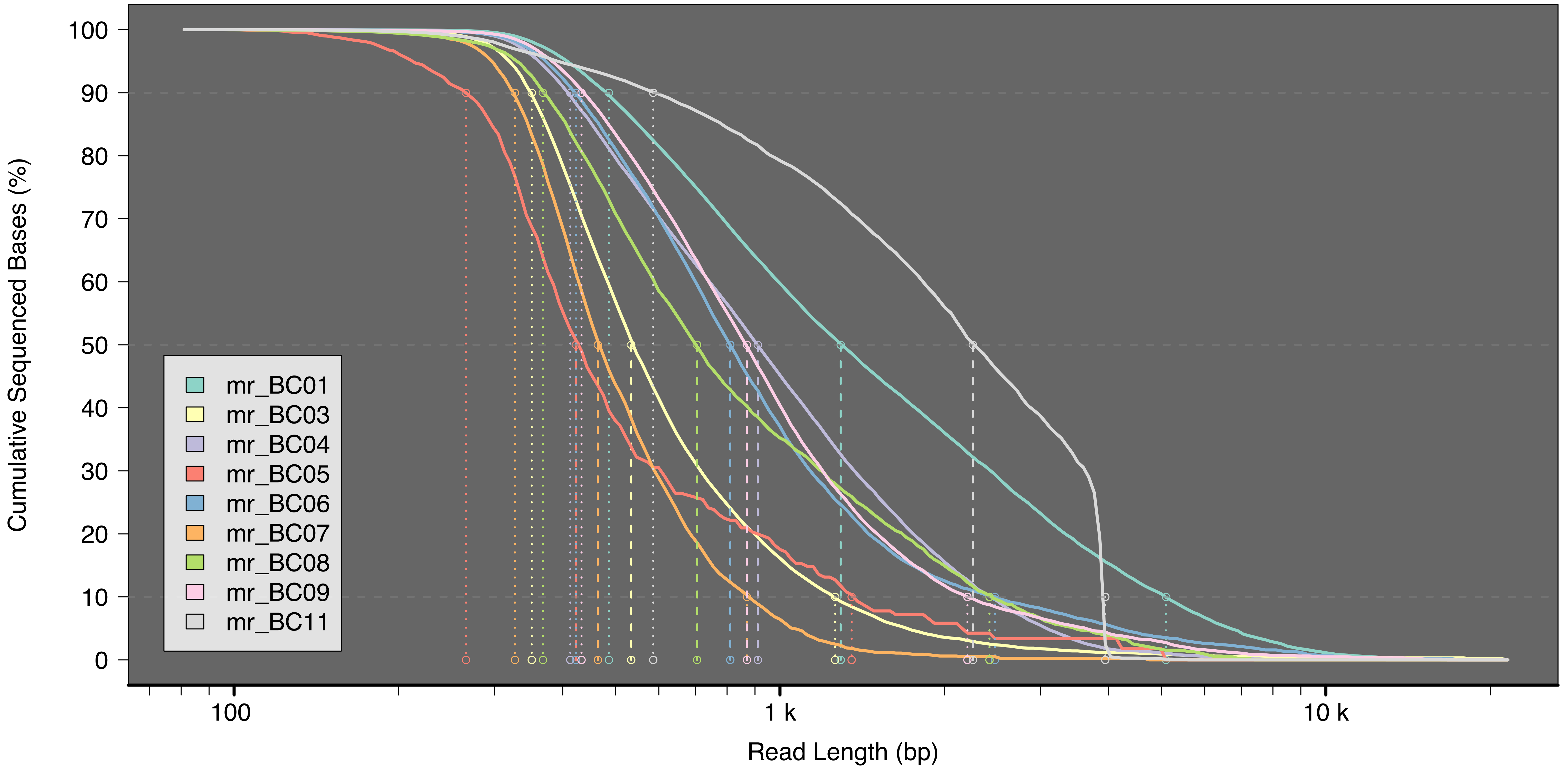

Cumulative Sequenced Bases (%)
100
90
80
70
60
50
40
30
20
10
0
mr_BC01
mr_BC03
mr_BC04
mr_BC05
mr_BC06
mr_BC07
mr_BC08
mr_BC09
mr_BC11
100
1 k
10 k
Read Length (bp)

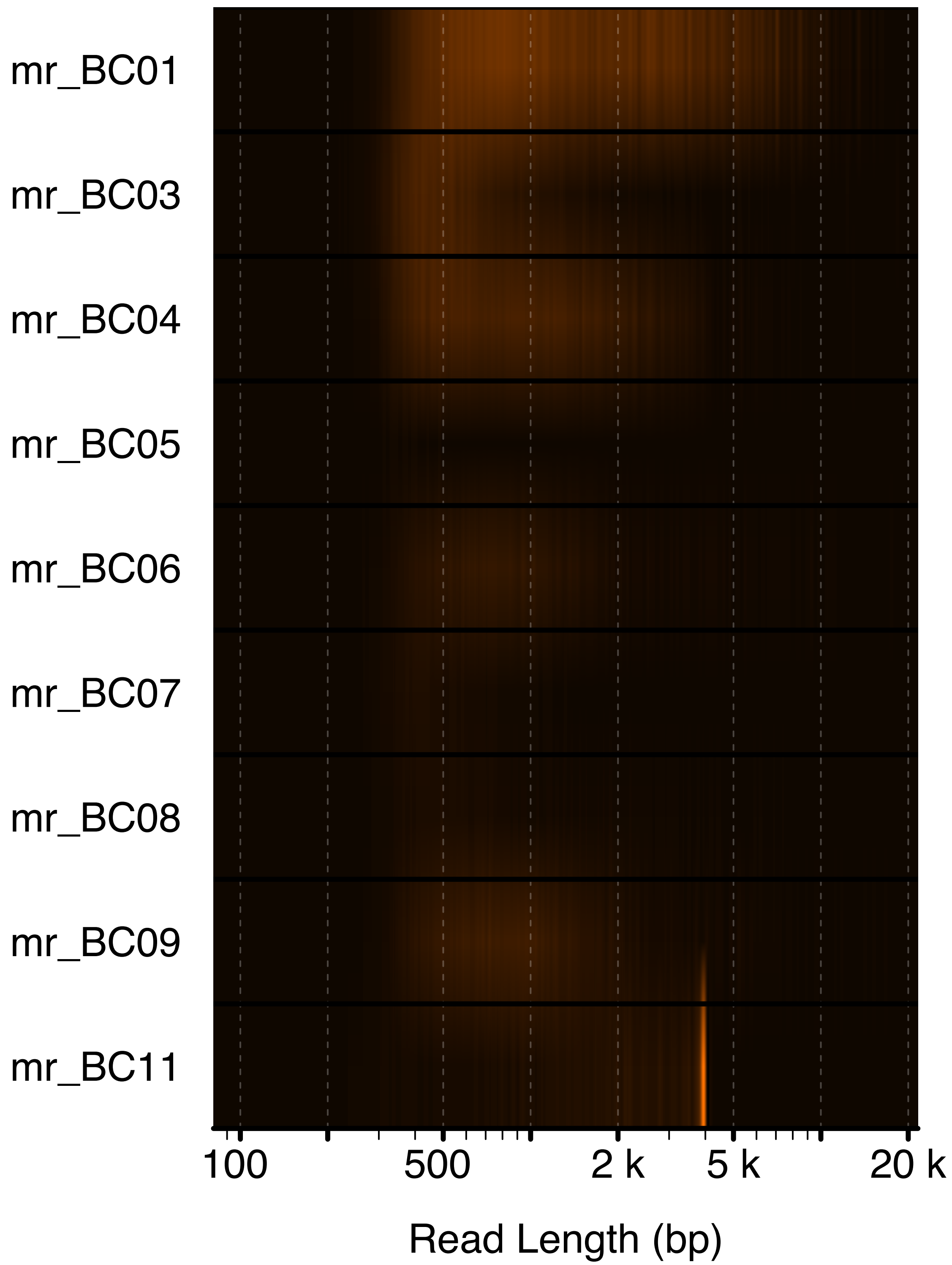
mr_BC01
mr_BC03
mr_BC04
mr_BC05
mr_BC06
mr_BC07
mr_BC08
mr_BC09
mr_BC11
100
500
2 k
5 k
20 k
Read Length (bp)

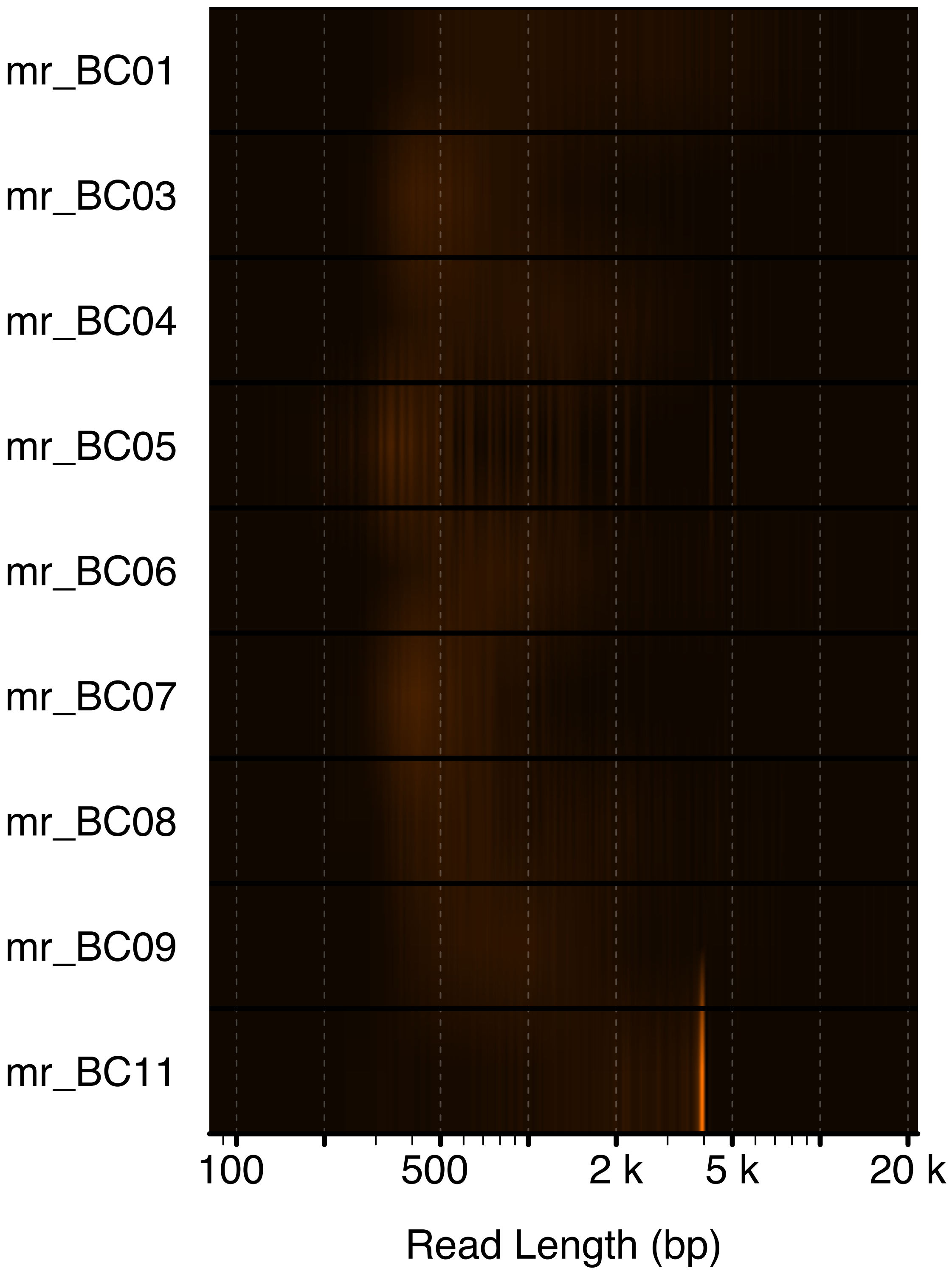
mr_BC01
mr_BC03
mr_BC04
mr_BC05
mr_BC06
mr_BC07
mr_BC08
mr_BC09
mr_BC11
100
500
2 k
5 k
20 k
Read Length (bp)

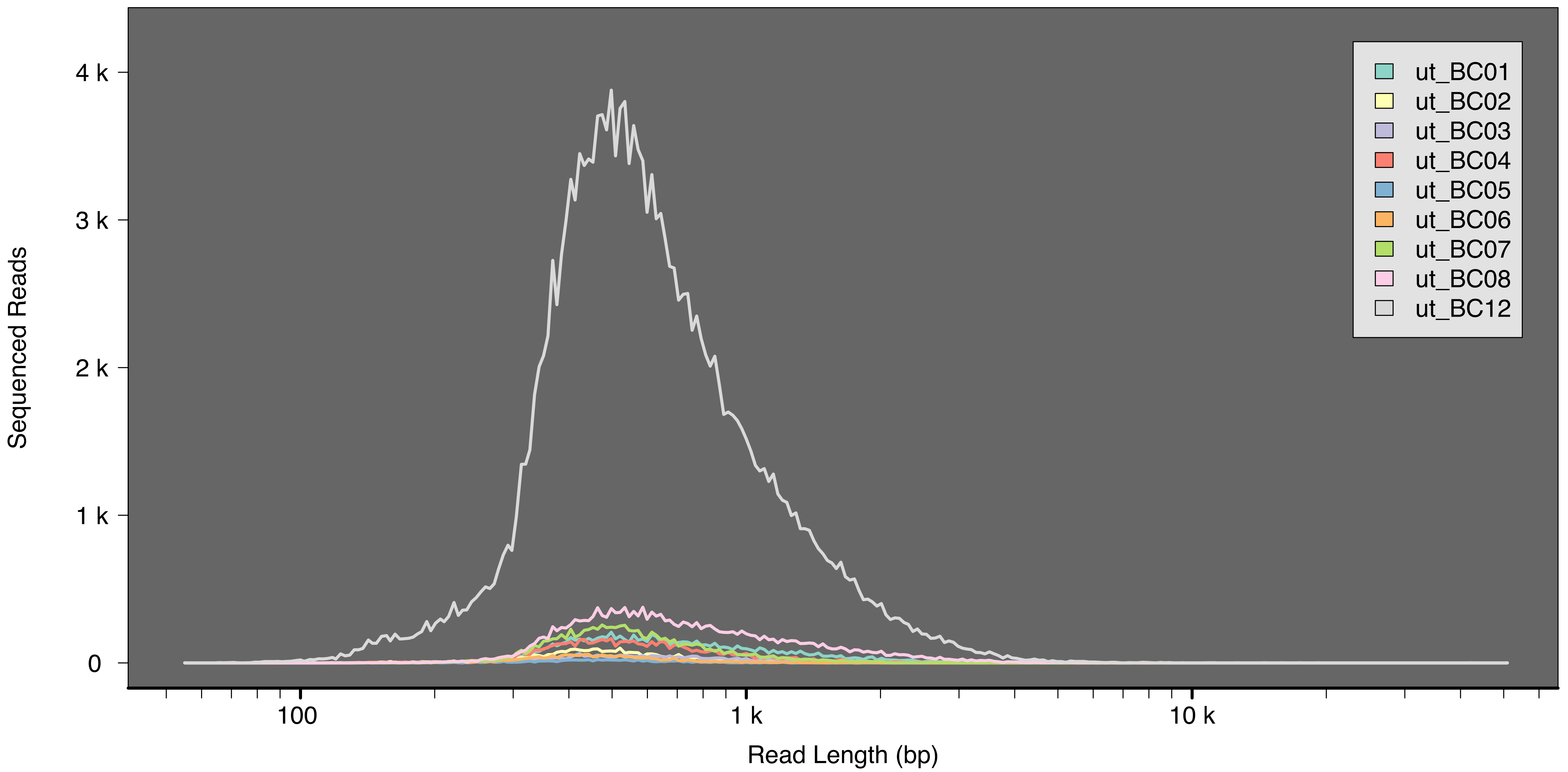

Sequenced Reads
4 k
3 k
2 k
1 k
0
100
1 k
10 k
Read Length (bp)
ut_BC01
ut_BC02
ut_BC03
ut_BC04
ut_BC05
ut_BC06
ut_BC07
ut_BC08
ut_BC12

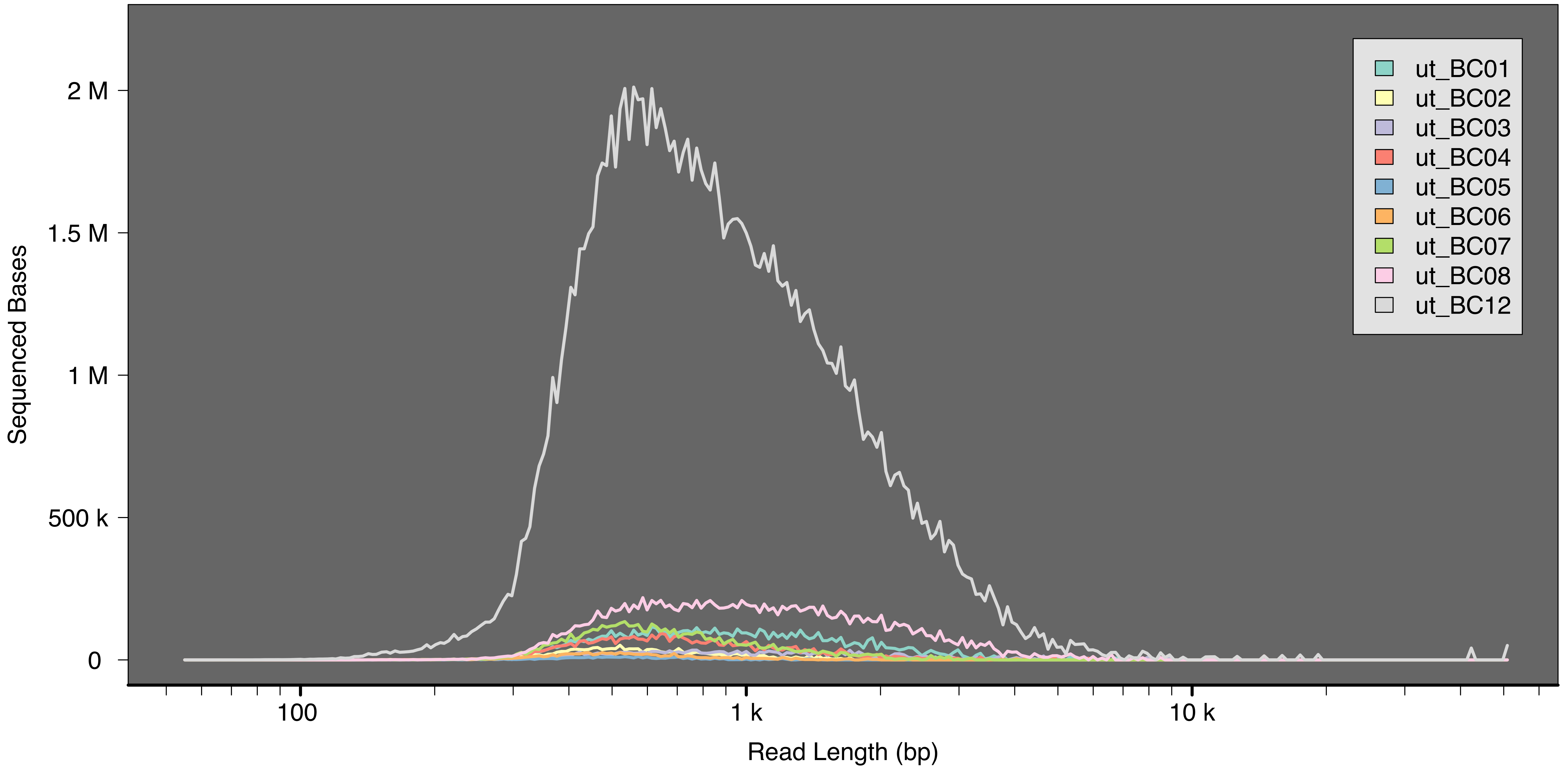

ut_BC01
ut_BC02
ut_BC03
ut_BC04
ut_BC05
ut_BC06
ut_BC07
ut_BC08
ut_BC12
Sequenced Bases
2 M
1.5 M
1 M
500 k
0
100
1 k
10 k
Read Length (bp)

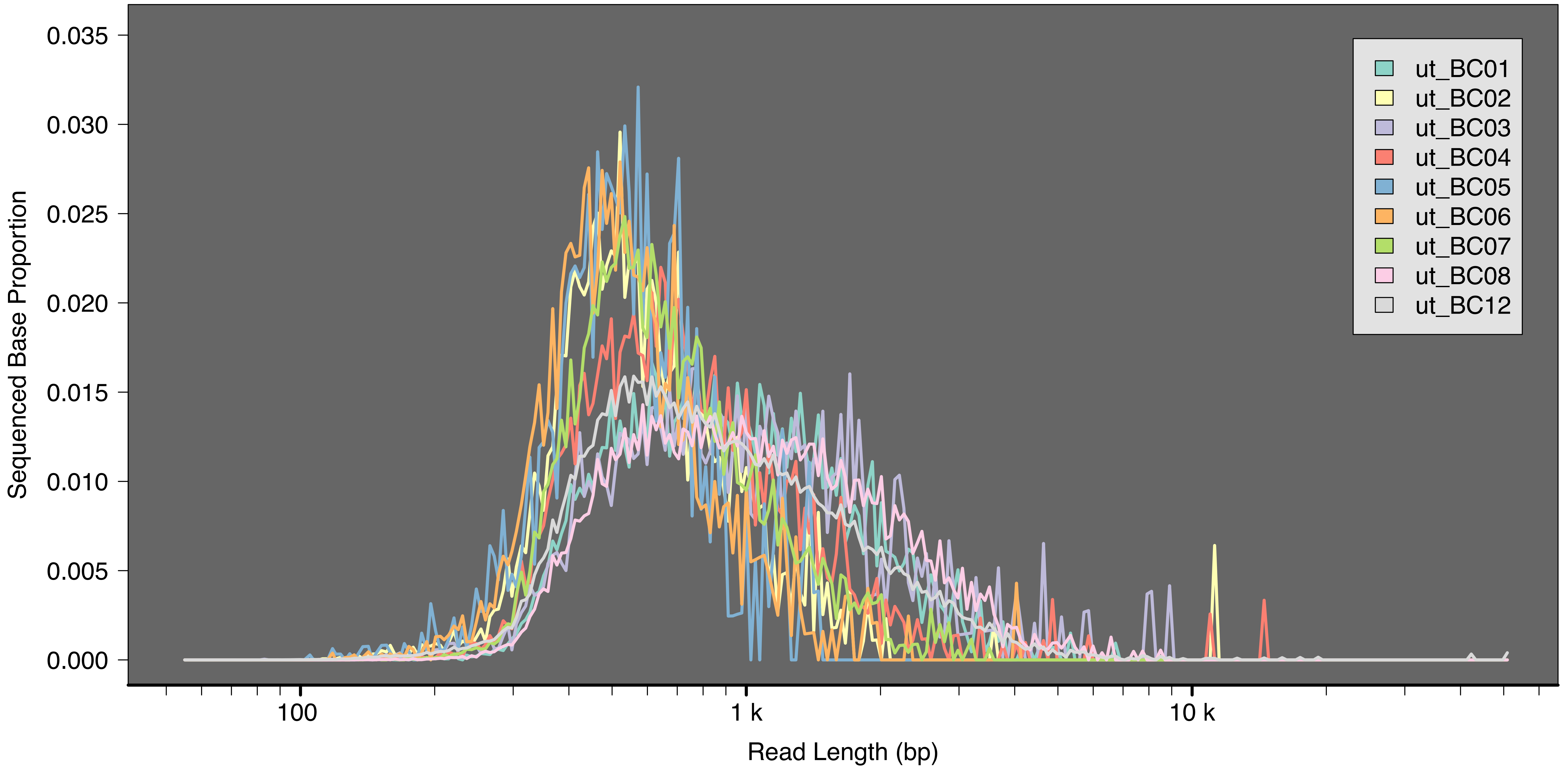

Sequenced Base Proportion
0.035
0.030
0.025
0.020
0.015
0.010
0.005
0.000
100
1 k
10 k
Read Length (bp)
ut_BC01
ut_BC02
ut_BC03
ut_BC04
ut_BC05
ut_BC06
ut_BC07
ut_BC08
ut_BC12

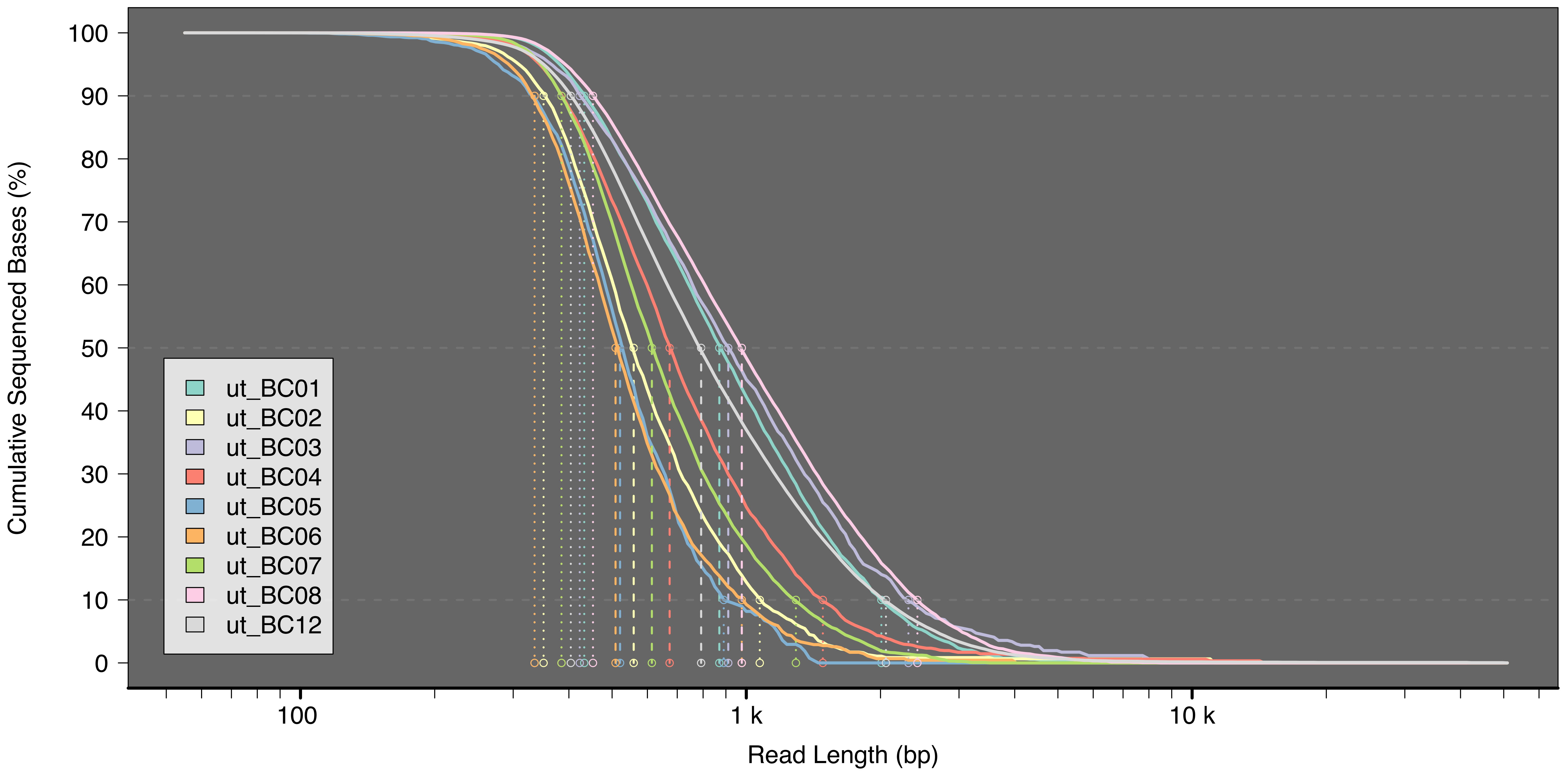
Cumulative Sequenced Bases (%)
100
90
80
70
60
50
40
30
20
10
0
ut_BC01
ut_BC02
ut_BC03
ut_BC04
ut_BC05
ut_BC06
ut_BC07
ut_BC08
ut_BC12
100
1 k
10 k
Read Length (bp)

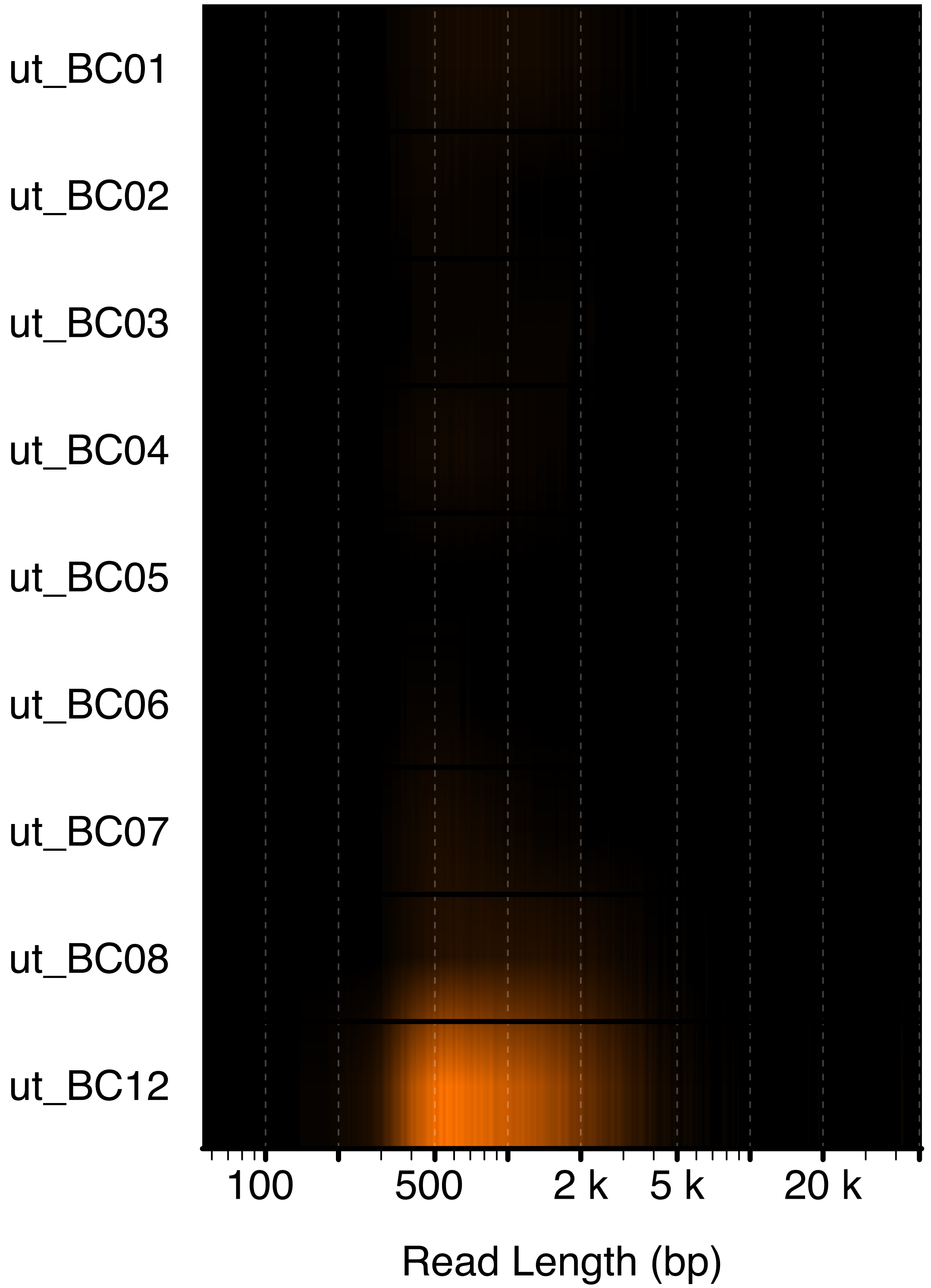

ut_BC01
ut_BC02
ut_BC03
ut_BC04
ut_BC05
ut_BC06
ut_BC07
ut_BC08
ut_BC12
100
500
2 k
5 k
20 k
Read Length (bp)

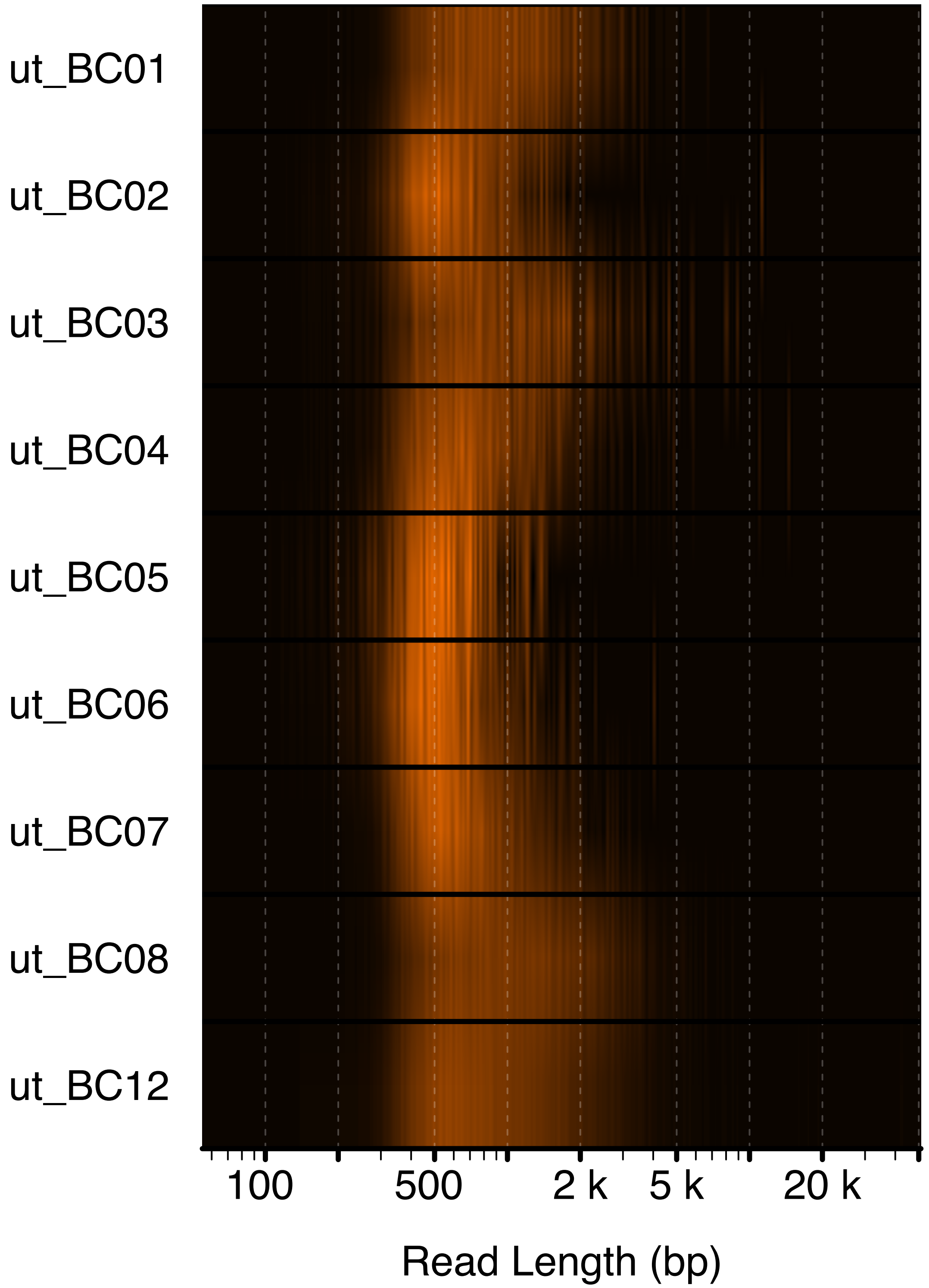
ut_BC01
ut_BC02
ut_BC03
ut_BC04
ut_BC05
ut_BC06
ut_BC07
ut_BC08
ut_BC12
100
500
2 k
5 k
20 k
Read Length (bp)

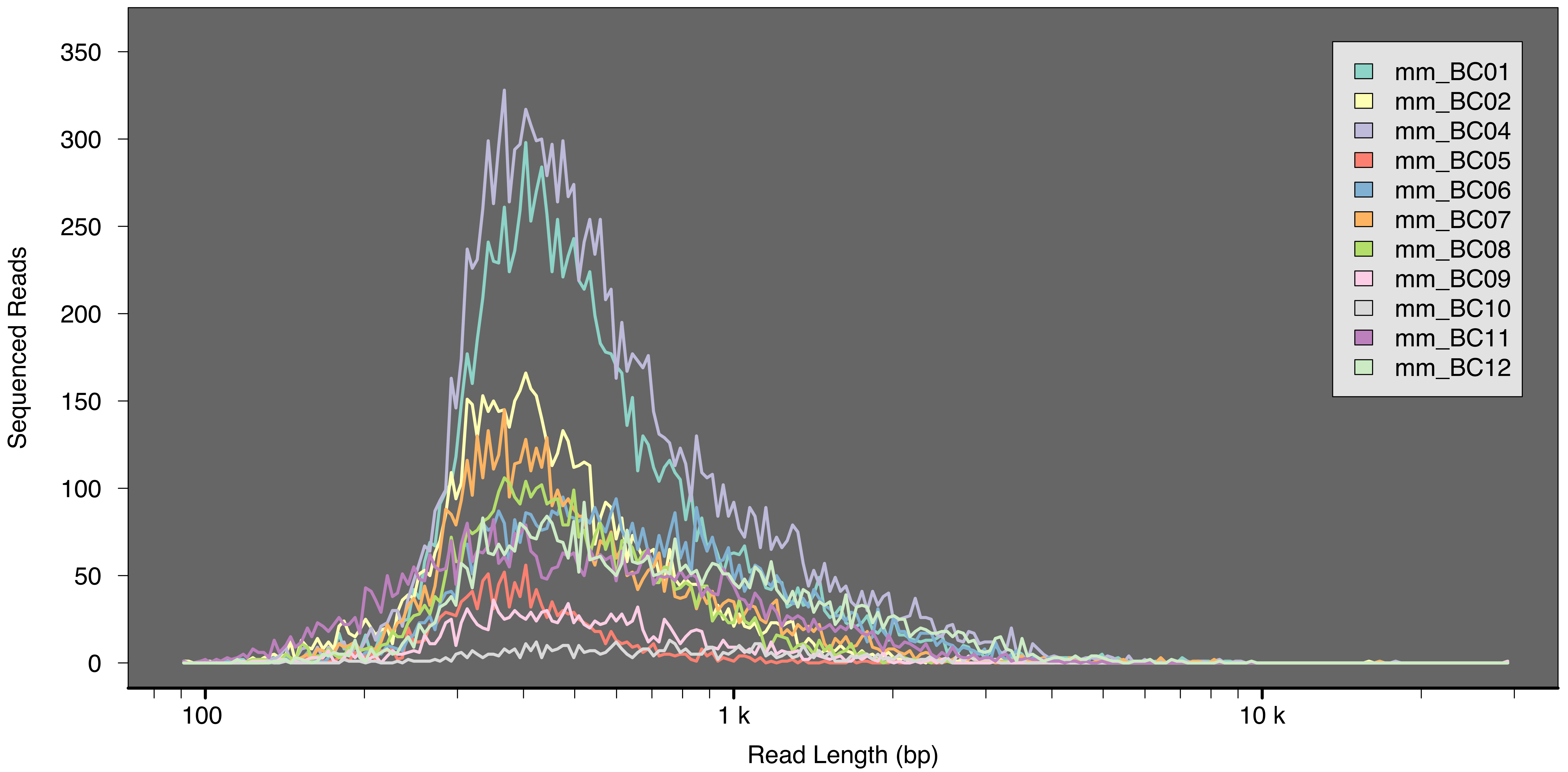

Sequenced Reads
Read Length (bp)
0
50
100
150
200
250
300
350
100
1 k
10 k
mm_BC01
mm_BC02
mm_BC04
mm_BC05
mm_BC06
mm_BC07
mm_BC08
mm_BC09
mm_BC10
mm_BC11
mm_BC12

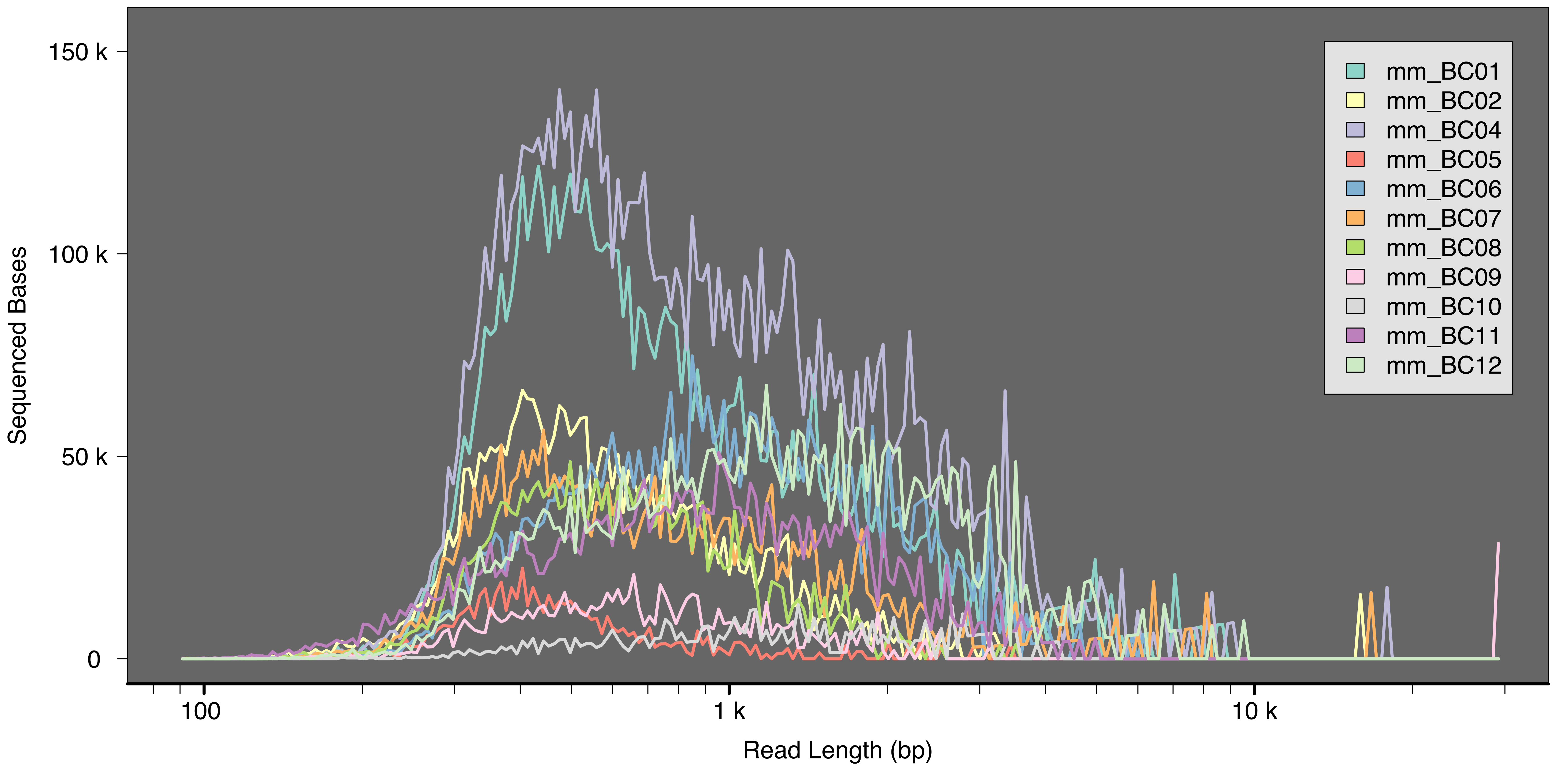
Sequenced Bases
150 k
100 k
50 k
0
100
1 k
10 k
Read Length (bp)
mm_BC01
mm_BC02
mm_BC04
mm_BC05
mm_BC06
mm_BC07
mm_BC08
mm_BC09
mm_BC10
mm_BC11
mm_BC12

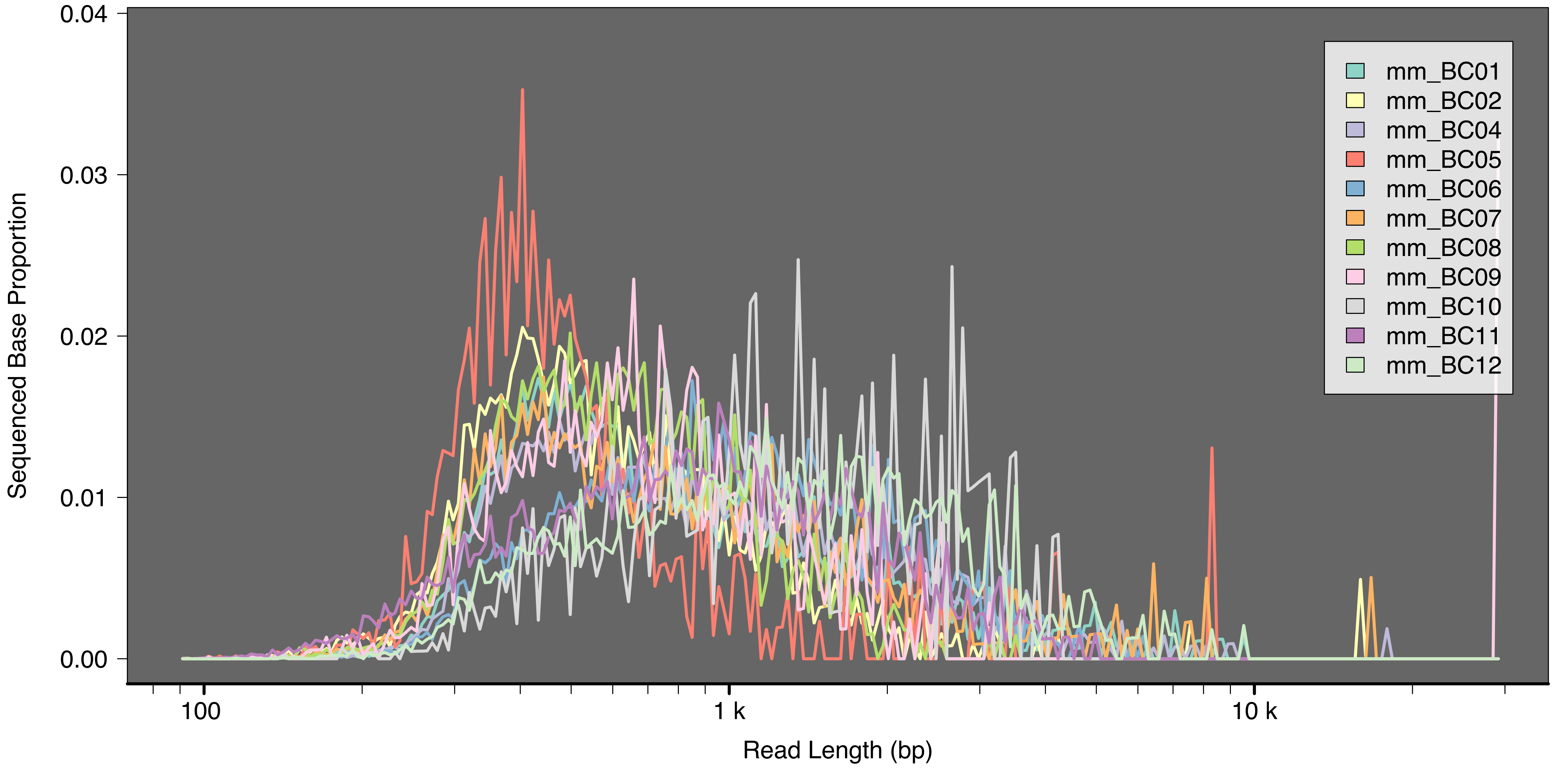
0.04
0.03
0.02
0.01
0.00
Sequenced Base Proportion
100
1 k
10 k
Read Length (bp)
mm_BC01
mm_BC02
mm_BC04
mm_BC05
mm_BC06
mm_BC07
mm_BC08
mm_BC09
mm_BC10
mm_BC11
mm_BC12

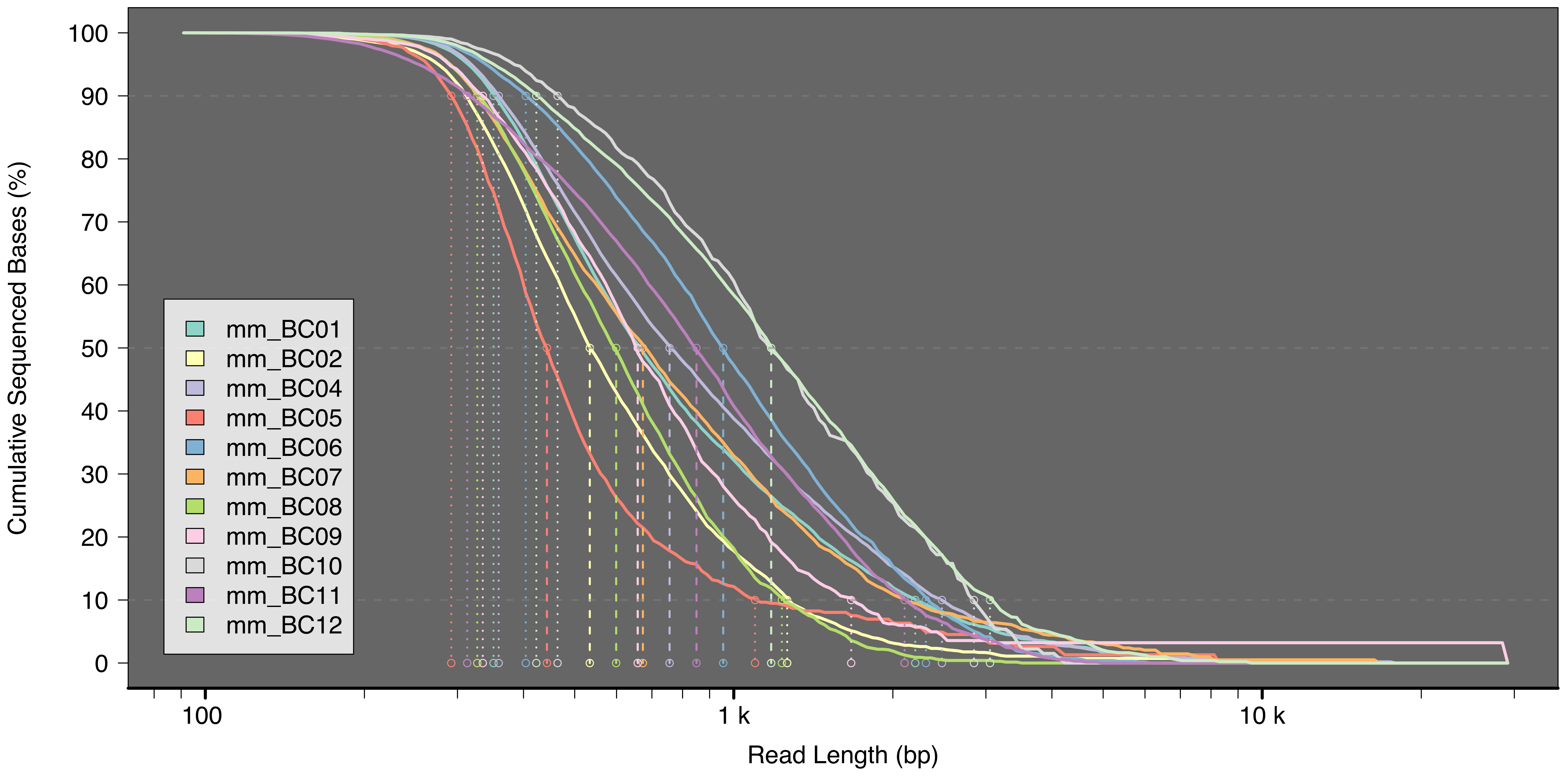

Cumulative Sequenced Bases (%)
100
90
80
70
60
50
40
30
20
10
0
Read Length (bp)
100
1 k
10 k
mm_BC01
mm_BC02
mm_BC04
mm_BC05
mm_BC06
mm_BC07
mm_BC08
mm_BC09
mm_BC10
mm_BC11
mm_BC12

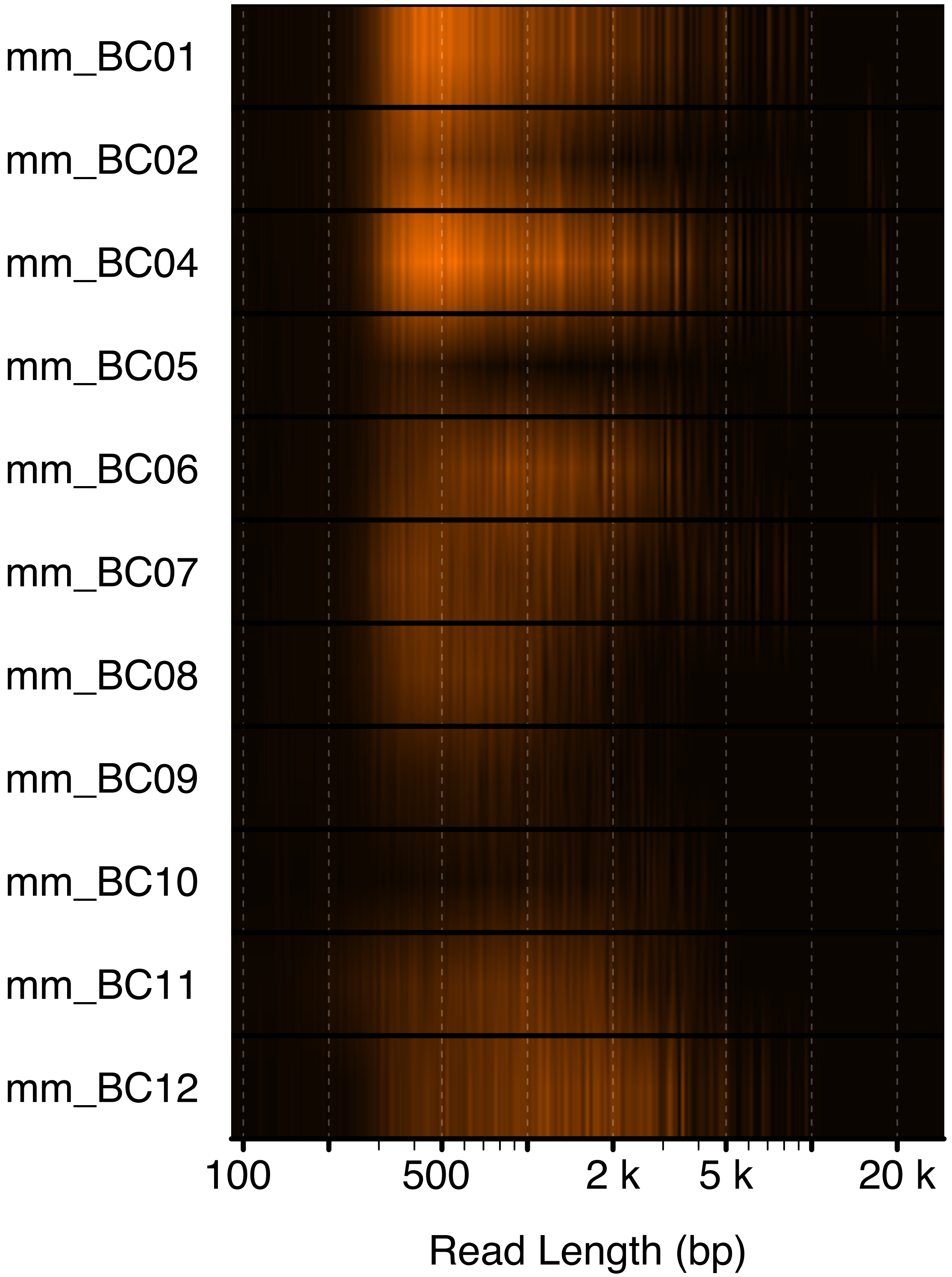
mm_BC01
mm_BC02
mm_BC04
mm_BC05
mm_BC06
mm_BC07
mm_BC08
mm_BC09
mm_BC10
mm_BC11
mm_BC12
100
500
2 k
5 k
20 k
Read Length (bp)

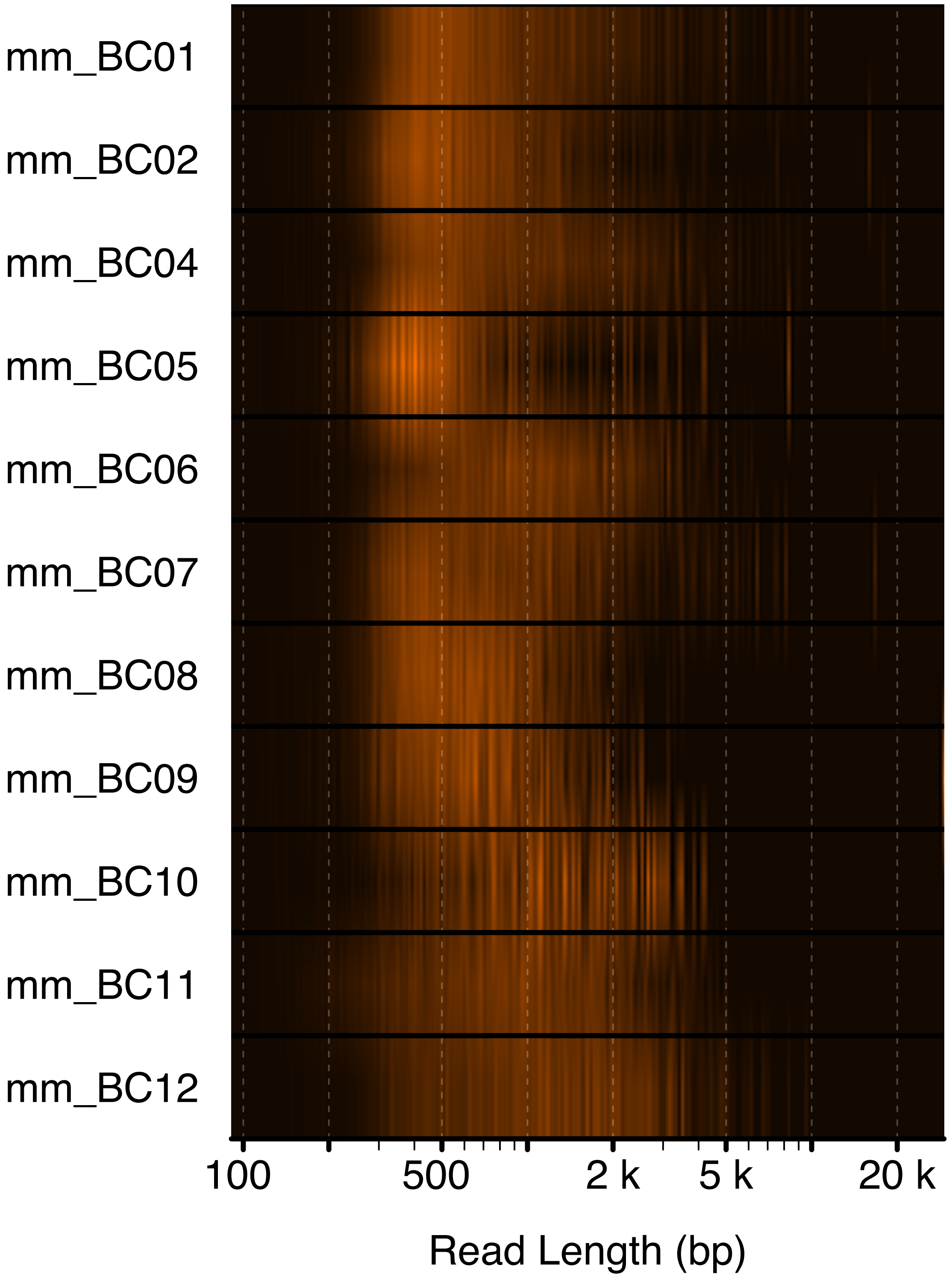

mm_BC01
mm_BC02
mm_BC04
mm_BC05
mm_BC06
mm_BC07
mm_BC08
mm_BC09
mm_BC10
mm_BC11
mm_BC12
100
500
2 k
5 k
20 k
Read Length (bp)
