## Supplemental File 2 for "Tree Lab: Portable genomics for early detection of plant viruses and pests in Sub-Saharan Africa"

Uploaded\_sample\_set\_\_-report.utf8.md


### Classification report for Uploaded sample set 1

###### *Pavian R package v0.8.3*

###### *Wed Jul 3 05:34:47 2019*

### Sample set summary

- Classification summary
- Raw read numbers
- Sample information

### Classification results

- Bacteria
- Viruses
- Eukaryotes
- Eukaryotes/Fungi
- Eukaryotes/Protists

Showing 100 of 7255 species.

Showing 100 of 358 species.

Showing 100 of 278 species.

```
## Warning in pavian:::build_sankey_network(my_report, nodePadding = 13,
## xScalingFactor = 0.9, : report does not contain any of the taxRanks -
## skipping it
```

### Sankey visualization

## mm\_BC01.

## mm\_BC02.

## mm\_BC03.

## mm\_BC04.

## mm\_BC05.

## mm\_BC06\_

## mm\_BC06.

## mm\_BC07.

## mm\_BC08.

## mm\_BC09.

## mm\_BC10.

## mm\_BC11.

## mm\_BC12.

## mr\_BC01.

## mr\_BC02.

## mr\_BC03.

## mr\_BC04.

## mr\_BC05.

## mr\_BC06.

## mr\_BC07.

## mr\_BC08.

## mr\_BC09.

## mr\_BC10.

## mr\_BC11.

## mr\_BC12.

## ut\_BC01.

## ut\_BC02.

## ut\_BC03.

## ut\_BC04.

## ut\_BC05.

## ut\_BC06.

## ut\_BC07.

## ut\_BC08.

## ut\_BC09.

## ut\_BC10.

## ut\_BC11.

## ut\_BC12.

### About

This file was generated with the Pavian R package version 0.8.3 on Wed Jul 3 05:35:01 2019. Please cite Pavian if you use it in your research.
